# Neutrophils promote CXCR3-dependent itch in the development of atopic dermatitis

**DOI:** 10.1101/653873

**Authors:** Carolyn M. Walsh, Rose Z. Hill, Jamie Schwendinger-Schreck, Jacques Deguine, Emily C. Brock, Natalie Kucirek, Ziad Rifi, Jessica Wei, Karsten Gronert, Rachel B. Brem, Gregory M. Barton, Diana M. Bautista

## Abstract

Chronic itch remains a highly prevalent disorder with limited treatment options. Most chronic itch diseases are thought to be driven by both the nervous and immune systems, but the fundamental molecular and cellular interactions that trigger the development of itch and the acute-to-chronic itch transition remain unknown. Here, we show that skin-infiltrating neutrophils are key initiators of itch in atopic dermatitis, the most prevalent chronic itch disorder. Neutrophil depletion significantly attenuated itch-evoked scratching in a mouse model of atopic dermatitis. Neutrophils were also required for several key hallmarks of chronic itch, including skin hyperinnervation, enhanced expression of itch signaling molecules, and upregulation of inflammatory cytokines, activity-induced genes, and markers of neuropathic itch. Finally, we demonstrate that neutrophils are required for induction of CXCL10, a ligand of the CXCR3 receptor that promotes itch via activation of sensory neurons, and we find that that CXCR3 antagonism attenuates chronic itch.

## Introduction

Chronic itch is a debilitating disorder that affects millions of people worldwide.^1–3^ It is a symptom of a number of skin diseases and systemic disorders, as well as a side effect of a growing list of medications. Like chronic pain, chronic itch can be a disease in and of itself.^4–6^ Unlike acute itch, which can facilitate removal of crawling insects, parasites, or irritants, persistent scratching in chronic itch disorders has no discernable benefit; scratching damages skin, leading to secondary infection, disfiguring lesions, and exacerbation of disease severity.^2, 7, 8^ The most common chronic itch disorder is atopic dermatitis (AD; commonly known as eczema), which affects fifteen million people in the United States alone.^9^ Severe AD can trigger the atopic march, where chronic itch and inflammation progress to food allergy, allergic rhinitis, and asthma.^9, 10^

Little is known about the underlying mechanisms that drive chronic itch pathogenesis. As such, studies of human chronic itch disorders have sought to identify candidate mechanisms of disease progression. A number of studies have identified biomarkers and disease genes in itchy human AD lesions.^11–15^ Indeed, a recent study compared the transcriptomes of healthy skin to itchy and non-itchy skin from psoriasis and AD patients, revealing dramatic changes in expression of genes associated with cytokines, immune cells, epithelial cells, and sensory neurons.^16^ However, due to the difficulty in staging lesion development and obtaining staged samples from patients, there is currently no temporal map of when individual molecules and cell types contribute to chronic itch pathogenesis. Furthermore, the use of human patient data does not allow for rigorous mechanistic study of how disease genes contribute to chronic itch. To this end, we used a well-characterized inducible animal model of itch to define where, when, and how these genes identified from patient data contribute to chronic itch pathogenesis.

We employed the MC903 mouse model of AD and the atopic march^17–21^ to provide a framework within which to identify the molecules and cells that initiate the development of atopic itch. The MC903 model is ideal for our approach because of its highly reproducible phenotypes that closely resemble human AD and the ability to induce the development of lesions and scratching.^18–20, 22–24^ By contrast, it is difficult to synchronously time the development of lesions in commonly used genetic models of AD, such as filaggrin mutant mice or Nc/Nga mice. Another advantage of the MC903 model is that it displays collectively more hallmarks of human AD than any one particular genetic mouse model. For example, the commonly used IL-31^tg^ overexpressor model^25, 26^ lacks strong Th2 induction,^27^ and itch behaviors have not yet been rigorously characterized in the keratinocyte-TSLP overexpressor model. As MC903 is widely used to study the chronic phase of AD, we hypothesized that MC903 could also be used to define the early mechanisms underlying the development of chronic itch, beginning with healthy skin. We performed RNA-seq of skin at key time points in the model. We complemented this approach with measurements of itch behavior and immune cell infiltration. The primary goal of our study was to identify the inciting molecules and cell types driving development of chronic itch. To that end, we show that infiltration of neutrophils into skin is required for development of chronic itch. Additionally, we demonstrate that neutrophils direct early hyperinnervation of skin, and the upregulation of itch signaling molecules and activity-induced genes in sensory neurons. Finally, we identify CXCL10/CXCR3 signaling as a key link between infiltrating neutrophils and sensory neurons that drives itch behaviors.

### MC903 triggers rapid changes in expression of skin barrier, epithelial cell-derived cytokine, and axon guidance genes

Although a variety of AD- and chronic itch-associated genes have been identified, when and how they contribute to disease pathogenesis is unclear. Using RNA-seq of MC903-treated skin, we observed distinct temporal patterns by which these classes of genes are differentially expressed across the first eight days of the model (Figure 1A-B, Figure 1-Figure Supplement 1A). Overall, we found that 62% of genes from a recent study of human chronic itch lesions^16^ (Figure 1-Figure Supplement 1A) and 67% of AD-related genes (Figure 1B) were significantly changed for at least one of the time points examined, suggesting that the MC903 mouse model recapitulates many key transcriptional changes occuring in human chronic itch and AD. MC903 dramatically alters the transcriptional profile of keratinocytes by derepressing genomic loci under the control of the Vitamin D Receptor. In line with rapid changes in transcription, proteases (*Klk6*, *Klk13*, among others) and skin barrier genes (*Cdhr1*) changed as early as six hours after the first treatment, before mice begin scratching (Figure 1B). Increased protease activity in AD skin is thought to promote breakdown of the epidermal barrier and release of inflammatory cytokines from keratinocytes.^28, 29^ One such cytokine, thymic stromal lymphopoetin (TSLP) is a key inducer of the Type 2 immune response, which is characteristic of human AD and the MC903 model, via signaling in CD4^+^ T cells, basophils, and other immune cells.^19, 20, 30–33^ Beginning at day two, before any significant itch-evoked scratching (Figure 1C), immune cell infiltration (Figure 1E-G, Figure 1-Figure Supplements 3A, 4A, 5A-C), or skin lesions (data not shown)^23^ were observed, we saw increases in *Tslp*, as well as several other epithelial-derived cytokines, including the neutrophil chemoattractant genes *Cxcl1, Cxcl2, Cxcl3,* and *Cxcl5* (Figure 1D). To ask whether upregulation of these chemokine genes was dependent on protease activity, we treated human keratinocytes with the Protease Activated Receptor 2 agonist SLIGRL. SLIGRL treatment triggered increased expression of several of these chemokine genes, including *IL8*, the human ortholog of mouse *Cxcl1/Cxcl2,* and *CXCL2* (Figure 1-Figure Supplement 6A). These increases occurred after a few hours of exposure to SLIGRL, suggesting that increased protease activity can rapidly trigger increases in neutrophil chemoattractants in skin, similar to what we observe in MC903-treated mouse skin.

**Figure 1.**
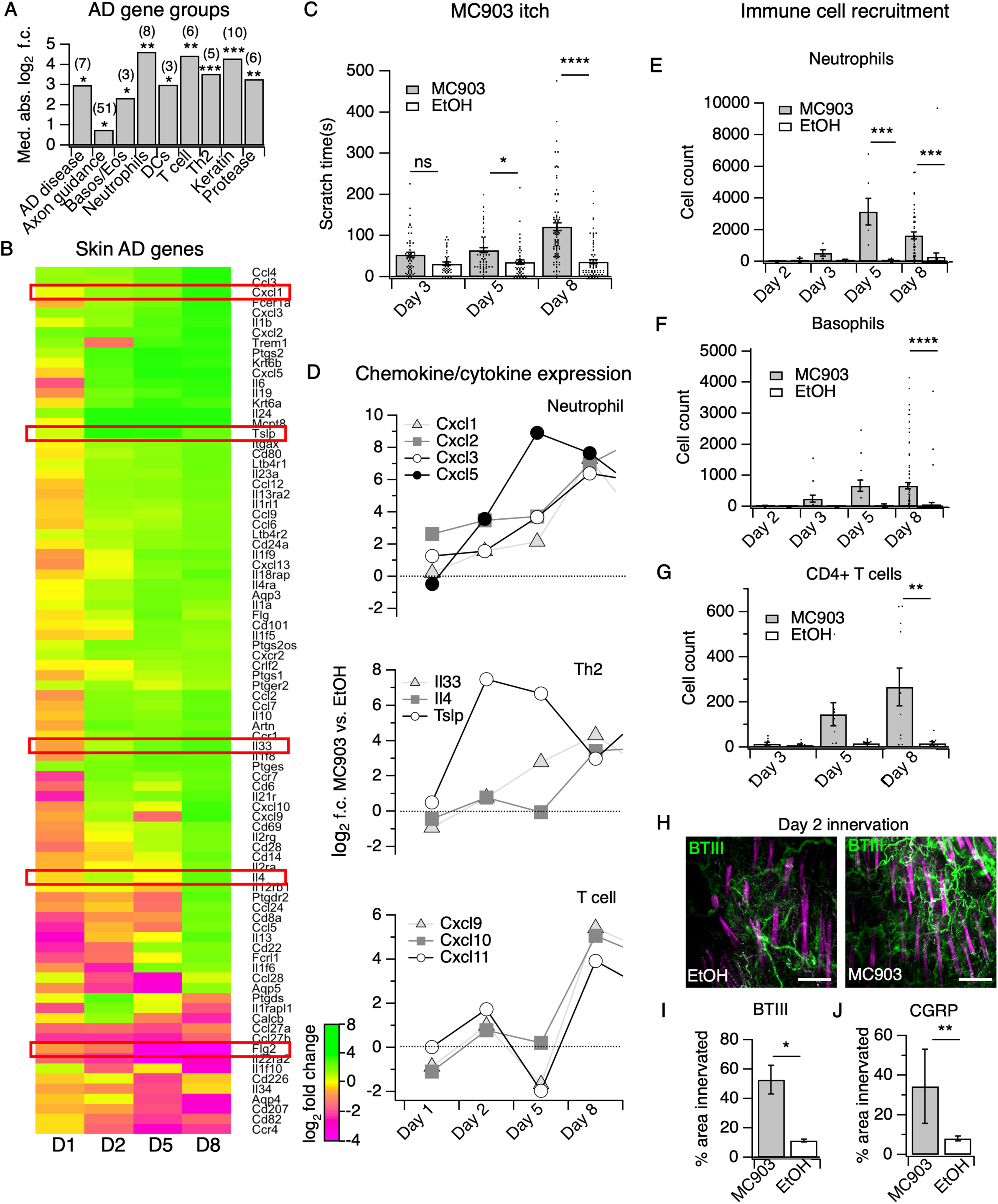
The MC903 model parallels the progression of human atopic disease and suggests a temporal sequence of AD pathogenesis. **A.** Exact permutation test (10,000 iterations, see Methods) for significance of mean absolute log_2_ fold change in gene expression at Day 8 (MC903 vs. ethanol) of custom-defined groups of genes for indicated categories (see **Figure 1-source data 1**). **B.** Log_2_ fold change in gene expression (MC903 vs. ethanol) in mouse skin at indicated time points for key immune and mouse/human AD genes that were significantly differentially expressed for at least one time point in the MC903 model. Only genes from our initial list (see Methods) differentially expressed at corrected *p* < 0.05 and changing > 2-fold between treatments for at least one condition are shown. Green bars = increased expression in MC903 relative to ethanol; magenta = decreased expression. Exact values and corrected *p*-values are reported in **Figure 1-source data 2** and **Supplemental Data**, respectively. D1 = 6 hours post-treatment; D2 = Day 2; D5 = Day 5; D8 = Day 8. **C.** Scratching behavior of mice treated with MC903 or ethanol for indicated length of time (two-way ANOVA: \*\*\*\**p*_interaction_< 0.0001, F(2,409) = 13.25; Sidak’s multiple comparisons: *p_day 3_* = 0.1309, n=62,51 mice; \**p_day 5_* = 0.0171, n=69,56 mice; \*\*\*\**p_day 8_* < 0.0001, n=92,85 mice). Exact values displayed in **Figure 1-source data 3**. **D.** Log_2_ fold change in gene expression of neutrophil chemoattractants (upper), Th2 cytokines (middle) and T cell chemoattractants (lower, from RNA-seq data). **E.** Neutrophil counts in MC903- and ethanol-treated skin at indicated time points (two-way ANOVA: \*\**p*_treatment_= 0.0023, F(1,102) = 9.82; Sidak’s multiple comparisons: *p_day 2_* > 0.999, n=4,4 mice; *p_day 3_* = 0.9801, n=5,5 mice; \*\*\**p_day 5_* = 0.0003, n=6,8 mice; \*\*\**p_day 8_* = 0.0001, n=40,38 mice). **F.** Basophil counts in MC903- and ethanol-treated skin at indicated time points (two-way ANOVA: \*\**p*_treatment_= 0.0051, F(1,102) = 8.17; Sidak’s multiple comparisons: *p_day 2_* > 0.999, n=4,4 mice; *p_day 3_* = 0.8850, n=5,5 mice; *p_day 5_* = 0.0606, n=6,8 mice; \*\*\*\**p_day 8_* < 0.0001, n=40,38 mice). **G.** CD4^+^ T cell counts in MC903- and ethanol-treated skin at indicated time points (two-way ANOVA: \*\**p*_time_= 0.0042, F(1,44) = 9.10; *p_day 3_* = 0.9998, n=8,6 mice; *p_day 5_* = 0.2223, n=9,8 mice; \*\**p_day 8_* = 0.0021, n=11,8 mice). Day 8 immune cell infiltrate represented as % of CD45^+^ cells in Figure 1-Figure Supplement 2A-B (see **Supplementary Table 3** for all experimental conditions). Exact values displayed in **Figure 1-source data 4** and representative FACS plots for myeloid and T cell gating shown in Figure 1-Figure Supplement 3A and Figure 1-Figure Supplement 4A. For Figure 4E-G, data from mice receiving i.p. injection of PBS (see Figure 4) in addition to MC903 or EtOH are also included. **H.** (Upper and Lower) Representative maximum intensity Z-projections from immunohistochemistry (IHC) of whole-mount mouse skin on Day 2 of the MC903 model. Skin was stained with neuronal marker beta-tubulin III (BTIII; green). Hair follicle autofluorescence is visible in the magenta channel. Images were acquired on a confocal using a 20x water objective. **I.** Quantification of innervation (see Methods) of mouse skin as determined from BTIII staining (*p = 0.012; two-tailed t-test (*t* =3.114; df =9); n = 7,4 images each from 2 mice per treatment). Day 1 IHC results as follows: 31.78 ± 18.39 % (MC903) and 31.51 ± 16.43 % (EtOH); p = 0.988; two-tailed unpaired t-test; n = 6 images each from 2 mice per treatment. Exact values are reported in **Figure 1-source data 5**. **J.** Quantification of CGRP^+^ nerve fibers (see Methods) in skin (\*\**p* = 0.0083; two-tailed t-test (*t* =2.868; df =25); n=15, 12 images from 3 mice per treatment). Exact values are reported in **Figure 1-source data 5**. Representative images in Figure 1-Figure Supplement 9A.

Unexpectedly, in the skin we observed early changes in a number of transcripts encoding neuronal outgrowth factors (*Ngf*, *Artn*) and axon pathfinding molecules (*Slit1, Sema3d, Sema3a*), some of which are directly implicated in chronic itch^34–38^; Figure 1-Figure Supplement 7A), prior to when mice began scratching. We thus used immunohistochemistry (IHC) of whole-mount skin to examine innervation at this time point. We saw increased innervation of lesions at day two but not day one of the model (Figure 1H-I, Figure 1-Figure Supplement 8A). Our RNA-seq data showed elevation in skin CGRP transcript, along with other markers of peptidergic nerve endings, specifically at day 2. Indeed, we saw an increase in CGRP^+^ innervation of skin at day 2 (Figure 1J, Figure 1-Figure Supplement 9A), which suggests that elevation of neuronal transcripts in skin is due to hyperinnervation of peptidergic itch and/or pain fibers. The increased innervation was surprising because such changes had previously only been reported in mature lesions from human chronic itch patients.^16, 39–44^ Our findings suggest that early hyperinnervation is promoted by local signaling in the skin and is independent of the itch-scratch cycle.

### Neutrophils are the first immune cells to infiltrate AD skin

By day five, mice exhibited robust itch behaviors (Figure 1C) and stark changes in a number of AD disease genes (Figure 1A-B). For example, loss-of-function mutations in filaggrin (*FLG)* are a major risk factor for human eczema.^45, 46^ Interestingly, *Flg2* levels sharply decreased at day five. In parallel, we saw continued and significant elevation in neutrophil and basophil chemoattractant genes (*Cxcl1,2,3,5*, and *Tslp*, Figure 1D). Using flow cytometry, we observed a number of infiltrating immune cells in the skin at day 5. Of these, we neutrophils were the most abundant immune cell subtype (Figure 1E, Figure 1-Figure Supplement 3A). It was not until day eight that we observed the classical AD-associated immune signature in the skin,^47^ with upregulation of *Il4, Il33* and other Th2-associated genes (Figure 1B, Figure 1D). We also observed increases in the T cell chemoattractant genes *Cxcl9, Cxcl10,* and *Cxcl11* (Figure 1D), which are thought to be hallmarks of chronic AD lesions in humans.^48, 49^ Neutrophils and a number of other immune cells that started to infiltrate on day five were robustly elevated in skin by day eight, including basophils (Figure 1F), CD4^+^ T cells (Figure 1G, Figure 1-Figure Supplement 4A), eosinophils (Figure 1-Figure Supplement 5C), and mast cells (Figure 1-Figure Supplement 5B), but not inflammatory monocytes (Figure 1-Figure Supplement 5A).

CD4^+^ T cells are ubiquitous in mature human AD lesions^50^ and promote chronic AD itch and inflammation. More specifically, they play a key role in IL4Rα-dependent sensitization of pruriceptors in the second week of the MC903 model.^22^ Thus, we were quite surprised to find that itch behaviors preceded significant CD4^+^ T cell infiltration. Therefore, neutrophils drew our attention as potential early mediators of MC903 itch. While neutrophil infiltration is a hallmark of acute inflammation, it remains unclear whether neutrophils contribute to the pathogenesis of chronic itch. Moreover, neutrophils release known pruritogens, including proteases, reactive oxygen species, and/or histamine, inflammatory lipids, and cytokines that sensitize and/or activate pruriceptors.^51, 52^ Increased levels of the prostaglandin PGE_2_ and the neutrophil-specific leukotriene LTB_4_ have also been reported in skin of AD patients.^53^ Indeed, by mass spectrometry, we observed increases in several of these inflammatory lipids, PGD_2_ and PGE_2_, as well as LTB_4_ and its precursor 5-HETE (Figure 1-Figure Supplement 10A) in MC903-treated skin, implicating neutrophils in driving AD itch and inflammation. Thus, we next tested the requirement of neutrophils to itch in the MC903 model.

### Neutrophils are required for early itch behaviors in the MC903 model of AD

We first asked whether neutrophils, the most abundant population of infiltrating immune cells in this chronic itch model, were required for MC903-evoked itch. Systemic depletion of neutrophils using daily injections of an anti-Gr1 (aGr1) antibody^54, 55^ dramatically attenuated itch-evoked scratching through the first eight days of the model (Figure 2A). Consistent with a key role for neutrophils in driving chronic itch, our depletion strategy significantly and selectively reduced circulating and skin infiltrating neutrophils on days five and eight, days on which control, but not depleted mice, scratched robustly (Figure 2B; Figure 2-Figure Supplement 1A-C). In contrast, basophils and CD4^+^ T cells continued to infiltrate the skin following aGr1 treatment (Figure 2C-D), suggesting that these cells are not required for early MC903 itch.

**Figure 2.**
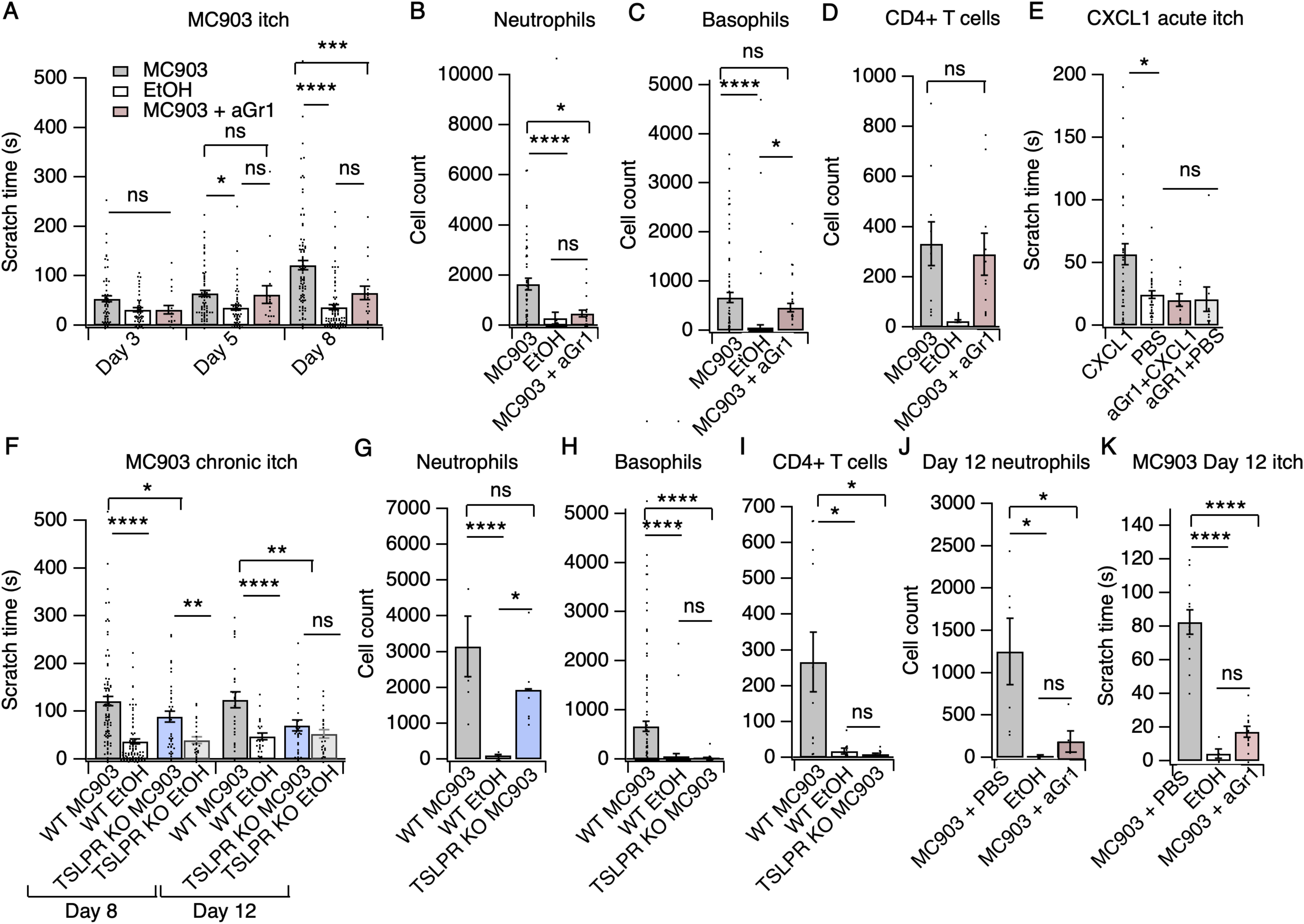
Neutrophils are necessary and sufficient for itch behaviors. **A.** Scratching behavior of uninjected and PBS-injected mice (combined) and aGr1-injected mice treated with MC903 or ethanol for indicated length of time (two-way ANOVA: \*\*\*\**p*_interaction_ < 0.0001, F(4,447) = 7.16; Tukey’s multiple comparisons: *p_day 3 MC903 vs. EtOH_* = 0.1111 n=62,51,17 mice; \**p_day 5 MC903 vs .EtOH_* = 0.0154, *p_day 5 MC903 vs. aGr1_* =0.9854, *p_day 5 aGr1 vs. EtOH_* =0.2267, n=69,56,17 mice; \*\*\*\**p_day 8 MC903 vs .EtOH_* < 0.0001, \*\*\**p_day 8 MC903 vs. aGr1_* = 0.0007, *p_day 8 aGr1 vs. EtOH_* = 0.1543, n=92,85,17 mice). **B.** Neutrophil count from cheek skin of uninjected/PBS-injected MC903- and ethanol-treated, and aGr1-injected MC903-treated mice on day 8 (one-way ANOVA: \*\*\*\**p* < 0.0001, F(2,92) = 10.59; Tukey’s multiple comparisons: \*\*\*\**p_MC903 vs. EtOH_* < 0.00001, n=40,38 mice; \**p_MC903 vs. aGr1 MC903_* = 0.0109, n=40,17 mice; *p_aGr1_* _vs. EtOH_ = 0.8859, n=38,17 mice). **C.** Basophil count from cheek skin of uninjected/PBS-injected MC903- and ethanol-treated, and aGr1-injected MC903-treated mice on day 8 (one-way ANOVA: \*\*\*\**p* = 0.0001, F(2,92) = 14.61; Tukey’s multiple comparisons: *p_MC903 vs. aGr1 MC903_* = 0.3217, n=40,17 mice, \*\*\*\**p_MC903 vs. EtOH_* < 0.0001, n=40,38 mice, \**p_aGr1 MC903 vs. EtOH_* = 0.0204, n=17,38 mice). **D.** CD4^+^ T cell count from cheek skin of PBS-injected MC903- and ethanol-treated, and aGr1-injected MC903-treated mice on day 8 (two-way ANOVA: \*\**p_treatment_* = 0.0035, F(1,35) = 9.82; Holm-Sidak multiple comparisons for PBS versus aGr1: *p_MC903_* = 0.8878, n=9,11 mice; *p_EtOH_* = 0.5201, n=8,9 mice). Control MC903 and EtOH data from **Figure 2B-C** are also displayed in Figure 1. Exact values displayed for **Figure 2A-D** in **Figure 2-source data 1**. **E.** Scratching behavior of mice immediately after injection of 1 µg CXCL1 or PBS (s.c. cheek). For neutrophil-depletion experiments, mice received 250 µg anti-Gr1 (aGr1) 20 hours prior to cheek injection of CXCL1 or PBS (one-way ANOVA: \*\*\*\**p* < 0.0001, F(4,88) = 75.53; Tukey’s multiple comparisons: \**p_CXCL1 vs. PBS_* = 0.0126, n=36,31 mice; *p_aGr1-CXCL1 vs. aGr1-PBS_* > 0.9999, n=10,10 mice; *p_aGr1-CXCL1_* _vs. PBS_ = 0.9986, n=10,31 mice). Exact values displayed in **Figure 2-source data 2**. **F.** Scratching behavior of WT and TSLPR^-/-^ (TSLPR KO) mice treated with MC903 or ethanol for indicated length of time (two-way ANOVA: \*\*\*\**p*_interaction_ < 0.0001, F(9,657) = 4.93; Tukey’s multiple comparisons: \*\*\*\**p_day 8 WT MC903 vs. EtOH_* < 0.0001, \**p_day 8 WT MC903 vs. KO MC903_* = 0.0194, *\*\*p_day 8 KO MC903 vs. KO EtOH_* = 0.0039, n=92,85,36,26 mice; \*\*\*\**p_day 12 WT MC903 vs. EtOH_* < 0.0001, \*\**p_day 12 WT MC903 vs. KO MC903_* = 0.0028, *p_day 12 KO MC903 vs. KO EtOH_* = 0.7061, n=26,26,27,23 mice). **G.** Neutrophil count from cheek skin of wild-type MC903- and ethanol-treated, and TSLPR^-/-^ MC903-treated mice on day 5 (two-way ANOVA: \*\**p*_genotype_ = 0.0025, F(2,125) = 6.28; Tukey’s multiple comparisons: \*\*\*\**p_day 5 WT MC903 vs. WT EtOH_* < 0.0001, n=6,8 mice; *p_day 5 WT MC903 vs. KO MC903_* = 0.2198, n=6,6 mice; \**p_day 5 WT EtOH vs. KO MC903_* = 0.0212, n=8,6 mice). **H.** Basophil count from cheek skin of wild-type MC903- and ethanol-treated, and TSLPR^-/-^ MC903-treated mice on day 8 (two-way ANOVA: \*\**p*_genotype_ = 0.0003, F(2,117) = 8.87; Tukey’s multiple comparisons: \*\*\*\**p_day 8 WT MC903 vs. WT EtOH_* < 0.0001, n=40,38 mice; \*\*\*\**p_day 8 WT MC903 vs. KO MC903_* < 0.0001, n=40,15 mice; *p_day 8 WT EtOH vs. KO MC903_* = 0.9519, n=38,15 mice). See also Figure 2-Figure Supplement 5A. For **Figures 2G-H**, data from days 3, 5, and 8 are presented in **Figure 2-source data 3. I.** CD4^+^ T cell count from cheek skin of wild-type MC903- and ethanol-treated, and TSLPR^-/-^ MC903-treated mice on day 8 (one-way ANOVA: \*\**p* = 0.0053, F(2,24) = 6.564; Tukey’s multiple comparisons: \**p_WT MC903 vs. WT EtOH_* = 0.0163, n=11,8 mice; \**p _MC903 vs. KO MC903_* = 0.0130, n=11,8 mice; *p_WT EtOH vs. KO MC903_* = 0.9953, n=8,8 mice). Wild-type MC903 and EtOH data from **2F-H** are also displayed in Figure 1. Exact values for **Figure 2F-I** displayed in **Figure 2-source data 3**. **J.** Neutrophil count from cheek skin of wild-type MC903- and ethanol-treated mice on day 12 of the MC903 model. MC903-treated animals received daily i.p. injections of 250 µg aGr1 antibody or PBS (250 µL) on days 8-11 of the model (one-way ANOVA: \**p* = 0.01, F(2,13) = 6.69; Tukey’s multiple comparisons: \**p_MC903-PBS vs. EtOH_* = 0.0141, n=6,5 mice; \**p_MC903-PBS vs. MC903-aGr1_* = 0.10330, n=6,5 mice; *p_MC903-aGr1 vs. EtOH_* = 0.9005, n=5,5 mice). **K.** Time spent scratching over a thirty minute interval for wild-type MC903- and ethanol-treated mice on day 12 of the MC903 model. MC903-treated animals received daily i.p. injections of 250 µg aGr1 antibody or PBS (250 µL) on days 8-11 of the model (one-way ANOVA: \*\*\*\**p* < 0.0001, F(2,26) = 53.1; Tukey’s multiple comparisons: \*\*\*\**p_MC903-PBS vs. EtOH_* < 0.0001, n=12,5 mice; \*\*\*\**p_MC903-PBS vs. MC903-aGr1_* < 0.0001, n=12,12 mice; *p_MC903-aGr1 vs. EtOH_* = 0.3734, n=12,5 mice). Values from bar plots are reported in **Figure 2-source data 5**.

We next used the cheek model of acute itch^56^ to ask whether neutrophil recruitment is sufficient to trigger scratching behaviors. As expected, we observed significant and selective recruitment of neutrophils to cheek skin within 15 minutes after CXCL1 injection (Figure 2-Figure Supplement 2A-B). CXCL1 injection also triggered robust scratching behaviors (Figure 2E) on a similar time course to neutrophil infiltration (Figure 2-Figure Supplement 2B). Thus, we next acutely depleted neutrophils with aGr1 to determine whether neutrophils were required for CXCL1-evoked acute itch. Indeed, aGr1-treatment rapidly reduced circulating neutrophils (Figure 2-Figure Supplement 2C) and resulted in a dramatic loss of CXCL1-evoked itch behaviors (Figure 2C). This effect was specific to neutrophil-induced itch, as injection of chloroquine, a pruritogen that directly activates pruriceptors to trigger itch, still triggered robust scratching in aGr1-treated animals (Figure 2-Figure Supplement 3A). Given that CXCL1 has been shown to directly excite and/or sensitize sensory neurons,^57, 58^ it is possible that the mechanism by which CXCL1 elicits itch may also involve neuronal pathways. However, our results show that CXCL1-mediated neutrophil infiltration is sufficient to drive acute itch behaviors, and that neutrophils are necessary for itch in the MC903 model.

We also examined MC903-evoked itch behaviors in mice deficient in *Crlf2*, the gene encoding the TSLP Receptor (TSLPR^-/-^ mice^59^). TSLPR is expressed by both immune cells and sensory neurons and is a key mediator of AD in humans and in mouse models.^18–20, 31, 60^ Surprisingly, MC903-treated TSLPR^-/-^ mice displayed robust scratching behaviors through the first eight days of the model (Figure 2F). In contrast to our results in aGr1-injected mice, TSLPR^-/-^ mice displayed robust neutrophil infiltration (Figure 2G), but completely lacked basophil and CD4^+^ T cell infiltration into the skin (Figure 2H-I, Figure 2-Figure Supplement 4A), and additionally displayed a reduction in mast cells (Figure 2-Figure Supplement 4A). These results suggest that basophils and CD4^+^ T cells are not required for early itch and further support an inciting role for neutrophils. Previous studies have shown that TSLP drives the expression of Type 2 cytokines and related immune cells that promote itch and inflammation in mature AD skin lesions.^18–20, 31, 60^ Consistent with a later role for TSLP signaling in AD, we did observe a significant reduction in itch-evoked scratching in TSLPR^-/-^ mice in the second week of the model (Figure 2F). Thus, our data support a model in which neutrophils are necessary for initiation of AD and itch behaviors early in the development of AD, whereas TSLPR signaling mediates the recruitment of basophils and CD4^+^ T cells to promote later stage itch and chronic inflammation.

The incomplete loss of itch behaviors on day 12 in the TSLPR^-/-^ animals (Figure 2F) raised the question of whether neutrophils might also contribute to itch during the second week of the MC903 model. To directly answer this question, we measured neutrophil infiltration and itch-evoked scratching on day 12 in mice that received either aGr1 or PBS on days 8-11 of the model to selectively deplete neutrophils solely during the second week. Neutrophil depletion in the second week with aGr1 robustly decreased skin-infiltrating neutrophils (Figure 2J), and substantially reduced scratching behaviors at day 12 (Figure 2K), supporting a role for neutrophils in chronic itch. Interestingly, we observed a 79% mean reduction in time spent scratching after neutrophil depletion at day 12, whereas loss of TSLPR effected a 44% reduction in time spent scratching. We speculate that neutrophils and TSLP signaling comprise independent mechanisms that together account for the majority of AD itch. In order to ascertain whether neutrophils could be salient players in other models of AD, and not just MC903, we measured neutrophil infiltration into ear skin in the 1-fluoro-2,4-dinitrobenzene (DNFB) model of atopic dermatitis, which relies on hapten-induced sensitization to drive increased IgE, mixed Th1/Th2 cytokine response, skin thickening, inflammation, and robust scratching behaviors in mice.^61–63^ Indeed, neutrophils also infiltrated DNFB- but not vehicle-treated skin (Figure 2-Figure Supplement 5A). Taken together, these observations are complementary to published datasets showing evidence for neutrophil chemokines and transcripts in human AD lesions.^11, 12, 13–15^ Overall, our data support a key role for neutrophils in promoting AD itch and inflammation.

### MC903 drives rapid and robust changes in the peripheral and central nervous systems

But how do neutrophils drive AD itch? Itchy stimuli are detected and transduced by specialized subsets of peripheral somatosensory neurons. Thus, to answer this question we first profiled the transcriptional changes in somatosensory neurons in the MC903 model, which were previously unstudied. In general, little is known regarding neuronal changes in chronic itch. Our initial examination of early hyperinnervation and changes in axon guidance molecules in skin suggested that neurons are indeed affected early on in the MC903 model, before the onset of itch-evoked scratching behaviors. In contrast to the skin, where we saw many early transcriptional changes, we did not see any significant transcriptional changes in the trigeminal ganglia (TG) until five days after the first treatment, and in total only 84 genes were differentially expressed through the eighth day (Figure 3A-B). These hits included genes related to excitability of itch sensory neurons,^51, 64^ neuroinflammatory genes,^65^ and activity-induced or immediate early genes (Figure 3A). Interestingly, we observed enrichment of neuronal markers expressed by one specific subset of somatosensory neurons that are dedicated to itch (*Il31ra, Osmr, Trpa1, Cysltr2,* and *Nppb*), termed “NP3” neurons.^51, 64, 66, 67^ Similar to what has been reported in mouse models of chronic pain, we observed changes in neuroinflammatory (*Bdnf, Nptx1, Nptx2, Nptxr*) and immune genes (*Itk, Cd19, Rag, Tmem173*). However, these transcriptional changes occurred just a few days after itch onset, in contrast to the slow changes in nerve injury and pain models that occur over weeks, indicating that neuropathic changes may occur sooner than previously thought in chronic itch. These changes occurred in tandem with the onset of scratching behaviors (Figure 1C), suggesting that the early molecular and cellular changes we observed by this time point may be important for development or maintenance of itch-evoked scratching.

**Figure 3.**
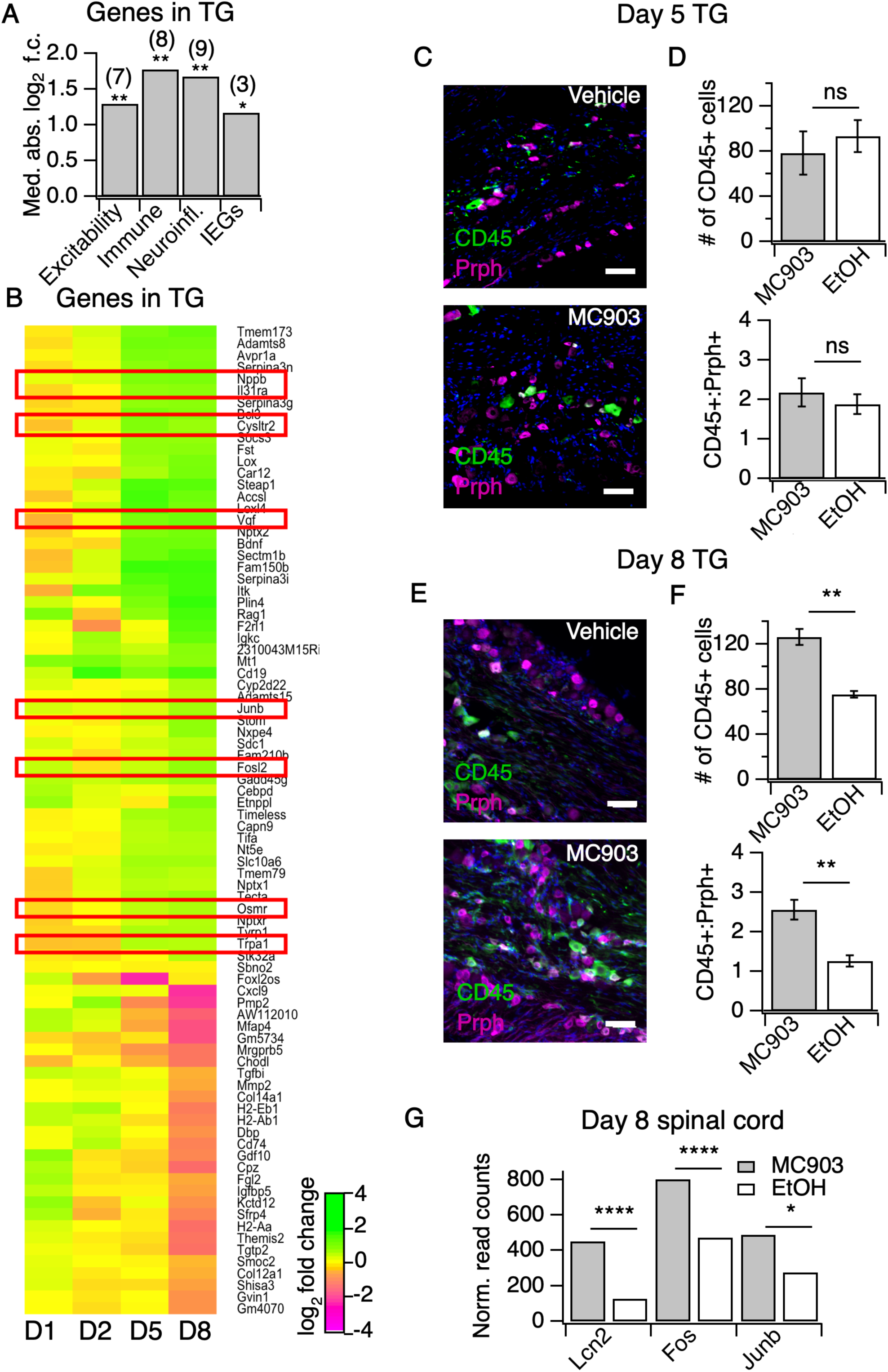
The MC903 model induces rapid and robust changes in neuronal tissue. **A.** Exact permutation test (10,000 iterations, see Methods) for significance of mean absolute log_2_ fold change in gene expression at Day 8 (MC903 vs. ethanol) of custom-defined groups of genes for indicated categories (see **Figure 3-source data 1**). **B.** Log_2_ fold change in gene expression (MC903 vs. ethanol) in mouse trigeminal ganglia (TG) at indicated time points for all genes which were significantly differentially expressed for at least one time point in the MC903 model. Green bars = increased expression in MC903 relative to ethanol; magenta = decreased expression. Exact values and corrected *p*-values are reported in **Figure 3-source data 2** and **Supplemental Data**, respectively. **C**. Representative composite images showing immune cells (CD45, green), and sensory neurons (Prph, magenta) with DAPI (blue) in sectioned trigeminal ganglia from mice treated with Vehicle or MC903 for five days on the cheek. **D**. Quantification of images examining average number of CD45^+^ cells per section and average ratio of CD45:Peripherin cells per section after five days of treatment (*p* = 0.562 (*t*=0.6318, *df*=4), 0.542 (*t*=0.6660, *df*=4); two-tailed unpaired t-tests, n=33-159 fields of view (images) each of both trigeminal ganglia from 3 mice per condition treated bilaterally). **E.** Representative composite images showing immune cells (CD45, green), and sensory neurons (Peripherin (Prph), magenta) with DAPI (blue) in sectioned trigeminal ganglia from mice treated with Vehicle or MC903 for eight days on the cheek. **F**. Quantification of images examining average number of CD45^+^ cells per section and average ratio of CD45:Peripherin cells per section after eight days of treatment (\*\**p* = 0.0019 (*t*=5.977,*df*=5), \*\**p =* 0.0093 (*t*=4.107,*df*=4); two-tailed unpaired t-tests; n=42-172 fields of view (images) each of both trigeminal ganglia from 3 EtOH or 4 MC903 animals treated bilaterally). Scale bar = 100 µm. Images were acquired on a fluorescence microscope using a 10x air objective. Values from bar plots and all TG IHC data are available in **Figure 3-source data 3**. **G.** Log_2_ fold change in gene expression (MC903 vs. ethanol) in mouse spinal cord on day 8 showing selected differentially expressed genes (*p_adjusted_ <* 0.05). Exact values and corrected *p*-values are reported in **Supplemental Data**.

The changes we observed in immune-related genes in the TG were suggestive of infiltration or expansion of immune cell populations, which has been reported in models of nerve injury and chronic pain, but has never been reported in chronic itch. To validate our observations, we used IHC to ask whether CD45^+^ immune cells increase in the TG. We observed a significant increase in TG immune cell counts at day eight but not day five (Figure 3C-F, Figure 3-Figure Supplement 1A-D). Because we observed such dramatic expression changes in the TG on day eight of the model, we postulated that the CNS may also be affected by this time point. Thus, we performed RNA-seq on spinal cord segments that innervate the MC903-treated rostral back skin of mice. To date, only one study has examined changes in the spinal cord during chronic itch.^68^ The authors showed that upregulation of the STAT3-dependent gene *Lcn2* occurred three weeks after induction of chronic itch and was essential for sustained scratching behaviors. Surprisingly, we saw upregulation of *Lcn2* on day eight of the MC903 model and, additionally, we observed robust induction of immediate early genes (*Fos, Junb,* Figure 3G), suggesting that MC903 itch drives activity-dependent changes in the spinal cord as early as one week after beginning treatment. Together, our findings show that sustained itch and inflammation can drive changes in the PNS and CNS much sooner than previously thought, within days rather than weeks after the onset of scratching. We next set out to explore how loss of neutrophils impacts the molecular changes observed in skin and sensory neurons in the MC903 model, and which of these changes might contribute to neutrophil-dependent itch.

### Neutrophils are required for upregulation of select itch- and atopic-related genes, including the itch-inducing chemokine CXCL10

To ask how neutrophils promote itch in the MC903 model, we examined the transcriptional changes in skin and sensory ganglia isolated from non-itchy neutrophil-depleted animals and from the TSLPR^-/-^ mice, which scratched robustly. A number of AD-associated cytokines that were upregulated in control MC903 skin were not upregulated in TSLPR^-/-^ and neutrophil-depleted skin. For example, *Il33* upregulation is both neutrophil- and TSLPR-dependent (Figure 4A, Figure 4-Figure Supplement 1A). By contrast, upregulation of epithelial-derived cytokines and chemokines *Tslp*, *Cxcl1*, *Cxcl2*, *Cxcl3*, and *Cxcl5* was unaffected by either loss of TSLPR or neutrophil depletion (Figure 4B), suggesting these molecules are produced by skin cells even when the MC903-evoked immune response is compromised. Consistent with previous studies, *Il4* upregulation was completely dependent on TSLPR but not neutrophils, establishing a role for TSLP signaling in the Type 2 immune response. Among the hundreds of MC903-dependent genes we examined, only a handful of genes were uniquely affected by neutrophil depletion. One such gene was *Cxcl10*, a chemokine known to be released by skin epithelial cells, neutrophils, and other myeloid cells.^52, 69–74^ *Cxcl10* expression was increased in TSLPR^-/-^ but not neutrophil-depleted skin (Figure 4B, Figure 4-Figure Supplement 1A). CXCL10 has been previously shown to drive acute itch in a model of allergic contact dermatitis via CXCR3 signaling in sensory neurons,^75^ and is elevated in skin of AD patients.^49^ Expression of *Cxcl9* and *Cxcl11*, two other CXCR3 ligands that are elevated in AD but have an unknown role in itch, was also decreased in AD skin of neutrophil-depleted mice (Figure 4B).

**Figure 4.**
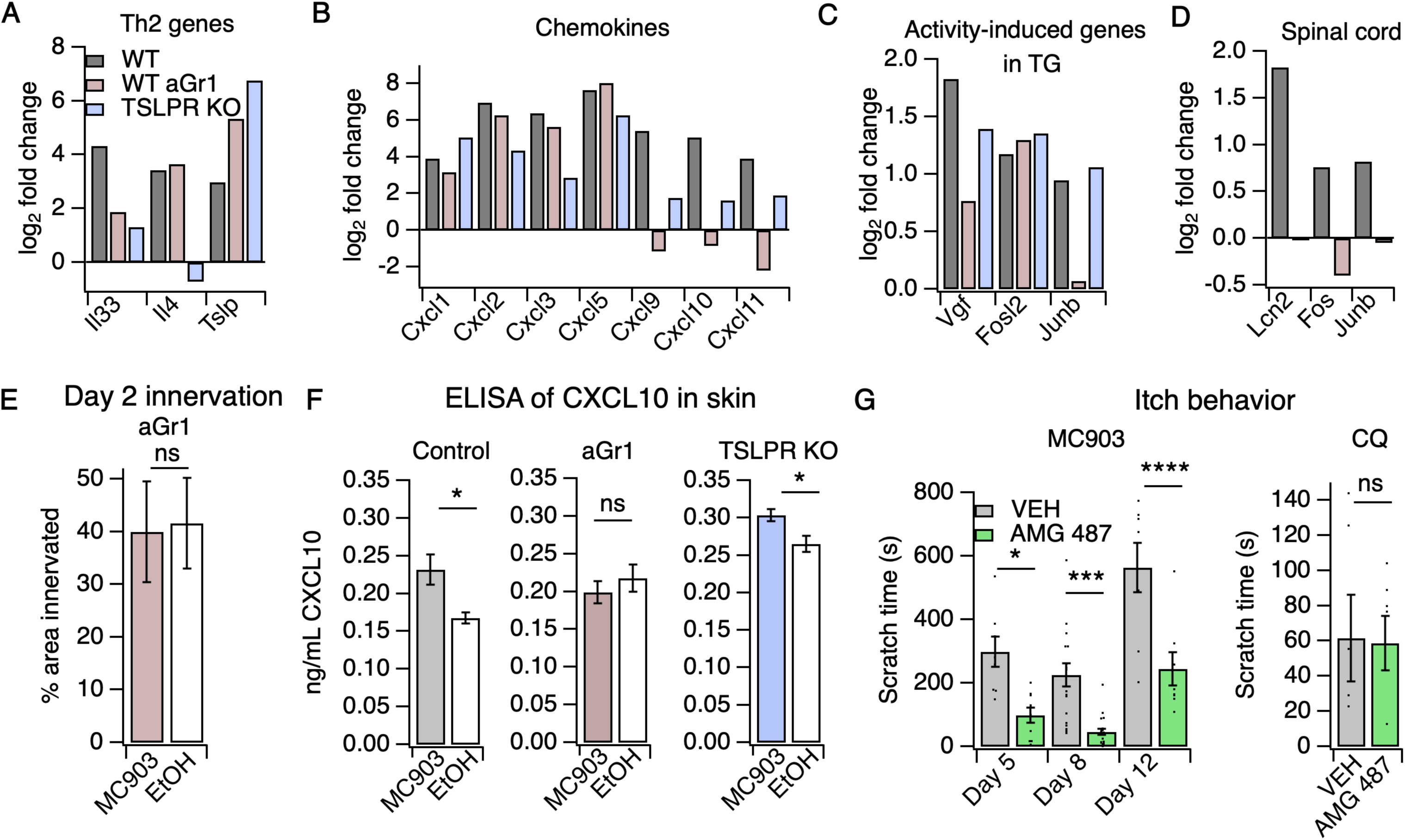
Neutrophils are required for induction of the itch-inducing chemokine CXCL10. **A.** Log_2_ fold change (Day 8 MC903 vs. EtOH) of Th2 genes in skin from uninjected wild-type, aGR1-treated, and TSLPR^-/-^ animals. **B.** Log_2_ fold change (Day 8 MC903 vs. EtOH) of chemokine genes in skin from uninjected wild-type, aGr1-treated, and TSLPR^-/-^ animals. **C.** Log_2_ fold change (Day 8 MC903 vs. EtOH) of activity-induced genes in trigeminal ganglia from uninjected wild-type, aGr1-treated, and TSLPR^-/-^ animals. **D.** Log_2_ fold change (Day 8 MC903 vs. EtOH) of *Lcn2* and activity-induced genes in spinal cord from uninjected and aGr1-treated wild-type mice on day 8. For **Figure 4A-D**, exact values and corrected *p*-values are reported in **Supplemental Data. E.** Quantification of innervation (see Methods) of MC903 and EtOH-treated mouse skin as determined from BTIII staining (*p* = 0.8985; two-tailed t-test (*t* =0.1294; df =18); n = 9,11 images each from 2 mice per treatment. Exact values are reported in **Figure 4-source data 1**. **F.** CXCL10 levels in skin homogenate as measured by ELISA on day 8 of the MC903 model for uninjected animals (left; \**p* = 0.029 (*t*=2.715, *df*=7); two-tailed t-test; n = 4,5 animals), animals which received aGr1 for 8 days (middle; *p* = 0.43 (*t*=0.815, *df*=11); two-tailed t-test; n = 6,6 animals), and TSLPR^-/-^ animals (right; \**p* = 0.0357 (*t*=2.696, *df*=6); two-tailed t-test; n = 4,4 animals. Skin homogenates were isolated on separate days and so uninjected, WT samples were not compared to aGr1-treated samples or to TSLPR^-/-^ samples. **G.** (Left) Time spent scratching over a thirty minute interval on days 5, 8, and 12 of the MC903 model, one hour after mice were injected with either 3.31 mM of the CXCR3 antagonist AMG 487 or vehicle (20% HPCD in PBS; 50 µL s.c. in rostral back); (two-way ANOVA: \*\*\*\**p*_treatment_ < 0.0001, F(1,67) = 50.64; Tukey’s multiple comparisons: \**p_day 5_* = 0.0216, n=8,10 mice; \*\*\**p_day 8_* = 0.0007, n=18,21 mice; \*\*\*\**p_day 12_* < 0.0001, n=8,8 mice). (Right) Time spent scratching over a thirty minute interval one hour after mice were injected with either 3.31 mM of the CXCR3 antagonist AMG 487 or vehicle (20% HPCD in PBS; 50 µL s.c. in rostral back), and immediately after mice were injected with 50 mM chloroquine (20 µL i.d., cheek). p = 0.92 (*t*=0.0964, *df*=8); two-tailed t-test; n = 5,5 mice. Values from bar plots in **Figures 4F-G** are displayed in **Figure 4-source data 2**.

### CXCR3 signaling is necessary for MC903-evoked chronic itch

We hypothesized that neutrophil-dependent upregulation of CXCL10 activates sensory neurons to drive itch behaviors. Consistent with this model, neutrophil depletion attenuated the expression of activity-induced immediate early genes (*Vgf, Junb*) in the TG, suggestive of neutrophil-dependent sensory neuronal activity (Figure 4C, Figure 4-Figure Supplement 1B). We found that neutrophils also contributed to other sensory neuronal phenotypes in the model. For example, we observed that expression of *Lcn2*, a marker of neuropathic itch, and activity-induced genes *Fos* and *Junb* were not increased in spinal cord isolated from neutrophil-depleted animals, indicating that neutrophil-dependent scratching behaviors may indeed drive changes in the CNS (Figure 4D). We also observed that neutrophil-depleted animals displayed no skin hyperinnervation at day two (Figure 4E). This result was surprising because we did not observe significant neutrophil infiltration at this early time point, but these data suggest that low numbers of skin neutrophils are sufficient to mediate these early effects.

To test our model wherein CXCL10 activates CXCR3 to drive neutrophil-dependent itch, we first asked whether this CXCR3 ligand is in fact released in MC903-treated skin. We performed ELISA on cheek skin homogenate and found that CXCL10 protein was increased in MC903-treated skin from uninjected wild-type and TSLPR^-/-^ animals, but not in skin from neutrophil-depleted mice (Figure 4F). To test whether CXCR3 signaling directly contributes to AD itch, we asked whether acute blockade of CXCR3 using the antagonist AMG 487^75^ affected scratching behaviors in the MC903 model. We found that the CXCR3 antagonist strongly attenuated scratching behaviors on days five, eight, and twelve (Figure 4G), with the greatest effect at day eight. In contrast, CXCR3 blockade did not attenuate scratching behaviors in naive mice injected with the pruritogen chloroquine (Figure 4G), demonstrating that CXCR3 signaling contributes to chronic itch but is not required for scratching in response to an acute pruritogen. Thus, we propose that neutrophils promote chronic itch in atopic dermatitis via upregulation of CXCL10 and subsequent activation of CXCR3-dependent itch pathways (Figure 5).

**Figure 5.**
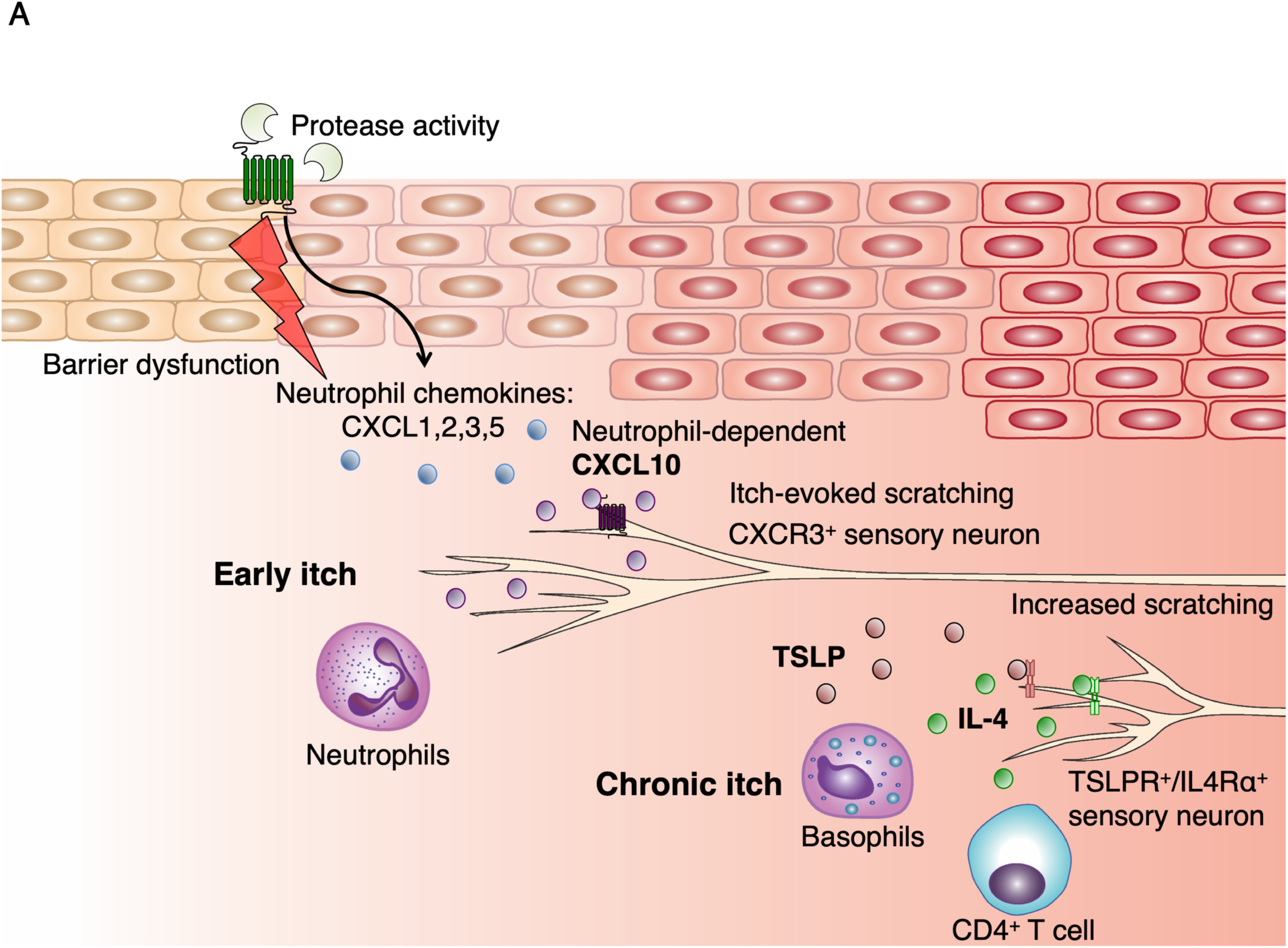
Model of early AD pathogenesis. **A.** AD induction first results in increased protease expression and barrier dysfunction, which drives production of the cytokines TSLP and CXCL1 via PAR2 activation within keratinocytes. CXCL1 can recruit neutrophils via its receptor CXCR2. Neutrophils may evoke itch by multiple pathways, including degranulation and release of proteases and histamine, production of sensitizing lipids such as PGE_2_ and LTB_4_,^52^ and induction of CXCL10 expression, which can activate sensory neurons via CXCR3. TSLP activates a number of immune cells to elicit IL-4 production, including basophils, which results in increased IL-4, recruitment of CD4^+^ T cells,^22^ and sensitization of neurons to promote itch later in the model.

## Discussion

There is great interest in unraveling the neuroimmune interactions that promote acute and chronic itch. Here, we show that neutrophils are essential for the early development of MC903-evoked itch. We further show that the recruitment of neutrophils to the skin is sufficient to drive itch behaviors within minutes of infiltration. While neutrophils are known to release a variety of pruritogens, their roles in itch and AD were not studied.^52^ Only a few studies have even reported the presence of neutrophils in human AD lesions.^12, 76–78^ Neutrophils have been implicated in psoriatic inflammation and inflammatory pain,^79–86^ where they are thought to rapidly respond to tissue injury and inflammation,^87^ but they have not been directly linked to itch.

There is a strong precedence for immune cell-neuronal interactions that drive modality-specific outcomes, such as itch versus pain, under distinct inflammatory conditions. In allergy, mast cells infiltrate the upper dermis and epidermis and release pruritogens to cause itch,^67, 88^ whereas in tissue injury, mast cell activation can trigger pain hypersensitivity.^89^ Likewise, neutrophils are also implicated in both pain and itch. For example, pyoderma gangrenosum, which causes painful skin ulcerations recruits neutrophils to the deep dermal layers to promote tissue damage and pain.^52^ In AD, neutrophils are recruited to the upper dermis and epidermis,^12, 78^ and we now show that neutrophils trigger itch in AD. Adding to the complex and diverse roles of neutrophils, neutrophils recruited to subcutaneous sites during invasive streptococcal infection alleviate pain by clearing the tissue of bacteria.^90^ Several potential mechanisms may explain these diverse effects of neutrophils. First, the location of the inflammatory insult could promote preferential engagement of pain versus itch nerve fibers.^52^ This is supported by observations that neutrophil-derived reactive oxygen species and leukotrienes can promote either itch or pain under different inflammatory conditions.^91–94^ Second, it has been proposed that there are distinct functional subsets of neutrophils that release modality-specific inflammatory mediators.^95^ Third, the disease-specific inflammatory milieu may induce neutrophils to specifically secrete mediators of either itch or pain. Indeed, all three of these mechanisms have been proposed to underlie the diverse functions of microglia and macrophages in homeostasis, tissue repair, injury, and neurodegenerative disease.^96^ It will be of great interest to the field to decipher the distinct mechanisms by which neutrophils and other immune cells interact with the nervous system to drive pain and itch.

In addition to neutrophils, TSLP signaling and the Type 2 immune response plays an important role in the development of itch in the second week of the MC903 model. Dendritic cells, mast cells, basophils, and CD4^+^ T cells are all major effectors of the TSLP inflammatory pathway in the skin. We propose that neutrophils play an early role in triggering itch and also contribute to chronic itch in parallel with the TSLP-Type 2 response. While we have ruled out an early role for TSLP signaling and basophils and CD4^+^ T cells in early itch, other cell types such as mast cells, which have recently been linked directly to chronic itch,^67, 88^ and dendritic cells may be playing an important role in setting the stage for itch and inflammation prior to infiltration of neutrophils.

Given the large magnitude of the itch deficit in the neutrophil-depleted mice, we were surprised to find fewer expression differences in MC903-dependent, AD-associated genes between neutrophil depleted and non-depleted mice than were observed between WT and TSLPR^-/-^ mice. One of the few exceptions were the Th1-associated genes *Cxcl9/10/11*.^11, 97^ We found that induction of these genes and of CXCL10 protein was completely dependent on neutrophils. While our results do not identify the particular cell type(s) responsible for neutrophil-dependent CXCL10 production, a number of cell types present in skin have been shown to produce CXCL10, including epithelial keratinocytes, myeloid cells, and sensory neurons.^52, 69–74^ In support of a role for neutrophils in promoting chronic itch, we observed striking differences in neutrophil-dependent gene expression in the spinal cord, where expression of activity-induced genes and the chronic itch gene *Lcn2* were markedly attenuated by loss of neutrophils. Moreover, we also demonstrate that depletion of neutrophils in the second week of the MC903 model can attenuate chronic itch-evoked scratching. In examining previous characterizations of both human and mouse models of AD and related chronic itch disorders, several studies report that neutrophils and/or neutrophil chemokines are indeed present in chronic lesions.^11–16, 98–102^ Our observations newly implicate neutrophils in setting the stage for the acute-to-chronic itch transition by triggering molecular changes necessary to develop a chronic, itchy lesion and also contributing to persistent itch.

Additionally, we demonstrate a novel role of CXCR3 signaling in MC903-induced itch. The CXCR3 ligand CXCL10 contributes to mouse models of acute and allergic itch;^75, 103, 104^ however, its role in chronic itch was previously unknown. We speculate that the residual itch behaviors after administration of the CXCR3 antagonist could be due to TSLPR-dependent IL-4 signaling, as TSLPR-deficient mice display reduced itch behaviors by the second week of the model, or due to some other aspect of neutrophil signaling, such as release of proteases, leukotrienes, prostaglandins, or reactive oxygen species, all of which can directly trigger itch via activation of somatosensory neurons.^52^ Our observations are in alignment with a recent study showing that dupilumab, a new AD drug that blocks IL4Rα, a major downstream effector of the TSLP signaling pathway, does not significantly reduce CXCL10 protein levels in human AD lesions.^105^ Taken together, these findings suggest that the TSLP/IL-4 and neutrophil/CXCL10 pathways are not highly interdependent, and supports our findings that *Il4* transcript is robustly upregulated in the absence of neutrophils. Additionally, targeting IL4Rα signaling has been successful in treating itch and inflammation in some, but not all, AD patients.^106^ We propose that biologics or compounds targeting neutrophils and/or the CXCR3 pathway may be useful for AD that is incompletely cleared by dupilumab monotherapy. Drugs targeting neutrophils are currently in clinical trials for the treatment of psoriasis, asthma, and other inflammatory disorders. For example, MDX-1100, a biologic that targets CXCL10, has already shown efficacy for treatment of rheumatoid arthritis in phase II clinical trials.^107^ While rheumatoid arthritis and AD have distinct etiologies,^108^ our body of work indicates that CXCL10 or CXCR3 may be promising targets for treating chronic itch. Our findings may also be applicable to other itch disorders where neutrophil chemoattractants and/or CXCL10 are also elevated, such as psoriasis and allergic contact dermatitis. Overall, our data suggest that neutrophils incite itch and inflammation in early AD through several mechanisms, including: 1) directly triggering itch upon infiltration into the skin, as shown by acute injection of CXCL1, and, 2) indirectly triggering itch by altering expression of endogenous pruritogens (e.g. induction of *Cxcl10* expression^52, 69–74^). Together, these direct and indirect mechanisms for neutrophil-dependent itch may explain why neutrophils have a dramatic effect on scratching behaviors on not only days eight and twelve but also day five of the model, when neutrophils are recruited in large numbers, but CXCR3 ligands are not as robustly induced.

More generally, our study provides a framework for understanding how and when human chronic itch disease genes contribute to the distinct stages of AD pathogenesis. Our analysis of MC903-evoked transcriptional changes suggests we may be able to extend findings in the model not only to atopic dermatitis, but also to related disorders, including specific genetic forms of atopy. For example, we provide evidence that MC903 treatment may also model the filaggrin loss-of-function mutations, which are a key inciting factor in human heritable atopic disease.^45, 46^ There are many rich datasets looking at mature patient lesions and datasets for mature lesions in other mouse models of chronic itch.^11–13, 15, 16, 22, 101, 109^ Our study adds a temporal frame of reference to these existing datasets and sets the stage for probing the function of AD disease genes in greater detail. Furthermore, we have mapped the time course of gene expression changes in primary sensory ganglia and spinal cord during chronic itch development. We show that the MC903 model recapitulates several hallmarks of neuropathic disease on a time course much shorter than has been reported for chronic itch, or chronic pain. Nervous system tissues are extremely difficult to obtain from human AD patients, and thus little is known regarding the neuronal changes in chronic itch disorders in both mouse models and human patients. Our findings can now be compared to existing and future datasets examining neuronal changes in chronic pain, diabetic neuropathy, shingles, neuropathic itch, psoriasis, and other inflammatory disorders where neuronal changes are poorly understood but may contribute to disease progression. The early changes we see in skin innervation, sensory ganglia, and spinal cord dovetail with recent studies examining neuroimmune interactions in other inflammatory conditions,^90, 110–112^ which all implicate early involvement of sensory neurons in the pathogenesis of inflammatory diseases.

**Figure 1-source data 1.** Values displayed in the bar plot shown in Figure 1A.

**Figure 1-source data 2.** Values displayed in the heat map shown in Figure 1B.

**Figure 1-source data 3**. Values displayed in the bar plot shown in Figure 1C.

**Figure 1-source data 4.** Values displayed in the bar plots shown in Figure 1E-G and Figure 1-Figure Supplement 5A-C.

**Figure 1-source data 5.** Values displayed in the bar plots shown in Figure 1I and Figure 1J.

**Figure 1-source data 6.** Values displayed in the heat map shown in Figure1-Figure Supplement 1A.

**Figure 1-source data 7.** Values displayed in the heat map shown in Figure1-Figure Supplement 6A.

**Figure 1-source data 8.** Values displayed in the heat map shown in Figure1-Figure Supplement 7A.

**Figure 1-source data 9.** Values displayed in the bar plot shown in Figure1-Figure Supplement 10A.

**Supplementary Table 1.** Number of mapped reads and sample information for all RNA-seq samples represented in the manuscript.

**Supplementary Table 2.** Outputs of statistical tests performed on behavioral and flow cytometry data to determine whether select data sets could be combined.

**Supplementary Table 3.** All flow cytometry data from Figures 1-2 represented as % of CD45^+^ cells.

**Supplemental Data.** DESeq differential expression output tables for all RNA-seq experiments in the manuscript.

**Figure 2-source data 1.** Values displayed in bar plots shown in Figure 2A-D.

**Figure 2-source data 2.** Values displayed in the bar plots shown in Figure 2E and Figs. 2-Figure Supplement 2-3.

**Figure 2-source data 3.** Values displayed in the bar plots shown in Figure 2F-I and Figure 2-Figure Supplement 4A-B.

**Figure 2-source data 4.** Values used to generate the line plots shown in Figure 2-Figure Supplement 1C.

**Figure 2-source data 5.** Values displayed in the bar plots shown in Figure 2J-K.

**Figure 2-source data 6.** Values displayed in the bar plots in Figure 2-Figure Supplement 5A.

**Figure 3-source data 1.** Values displayed in the bar plot shown in Figure 3A.

**Figure 3-source data 2.** Values displayed in the heat map shown in Figure 3B.

**Figure 3-source data 3.** Quantification of all IHC samples from trigeminal ganglia, and Values displayed in the bar plots shown in Figure 3D,F.

**Figure 4-source data 1.** Values displayed in the bar plot shown in Figure 4E.

**Figure 4-source data 2.** Values displayed in the bar plots shown in Figure 4F-G.

**Figure 4-source data 3.** Values displayed in the heat map shown in Figure 4-Figure Supplement 1A.

**Figure 4-source data 4.** Values displayed in the heat map shown in Figure 4-Figure Supplement 1B.

**Supplementary Data** - The outputs of all differential expression analyses used to determine adjusted *p* value and log_2_ fold change for RNA-seq experiments.

## Methods

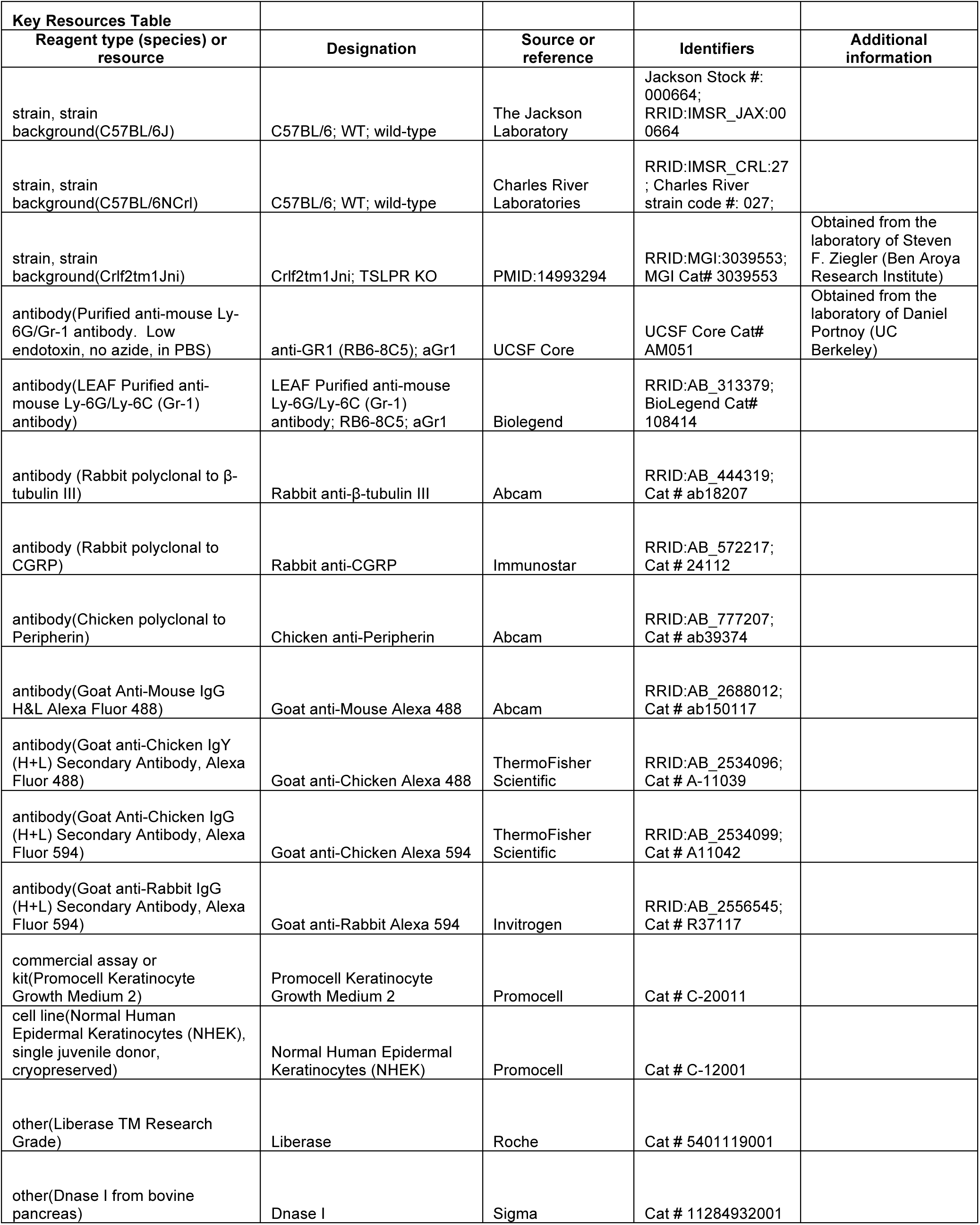
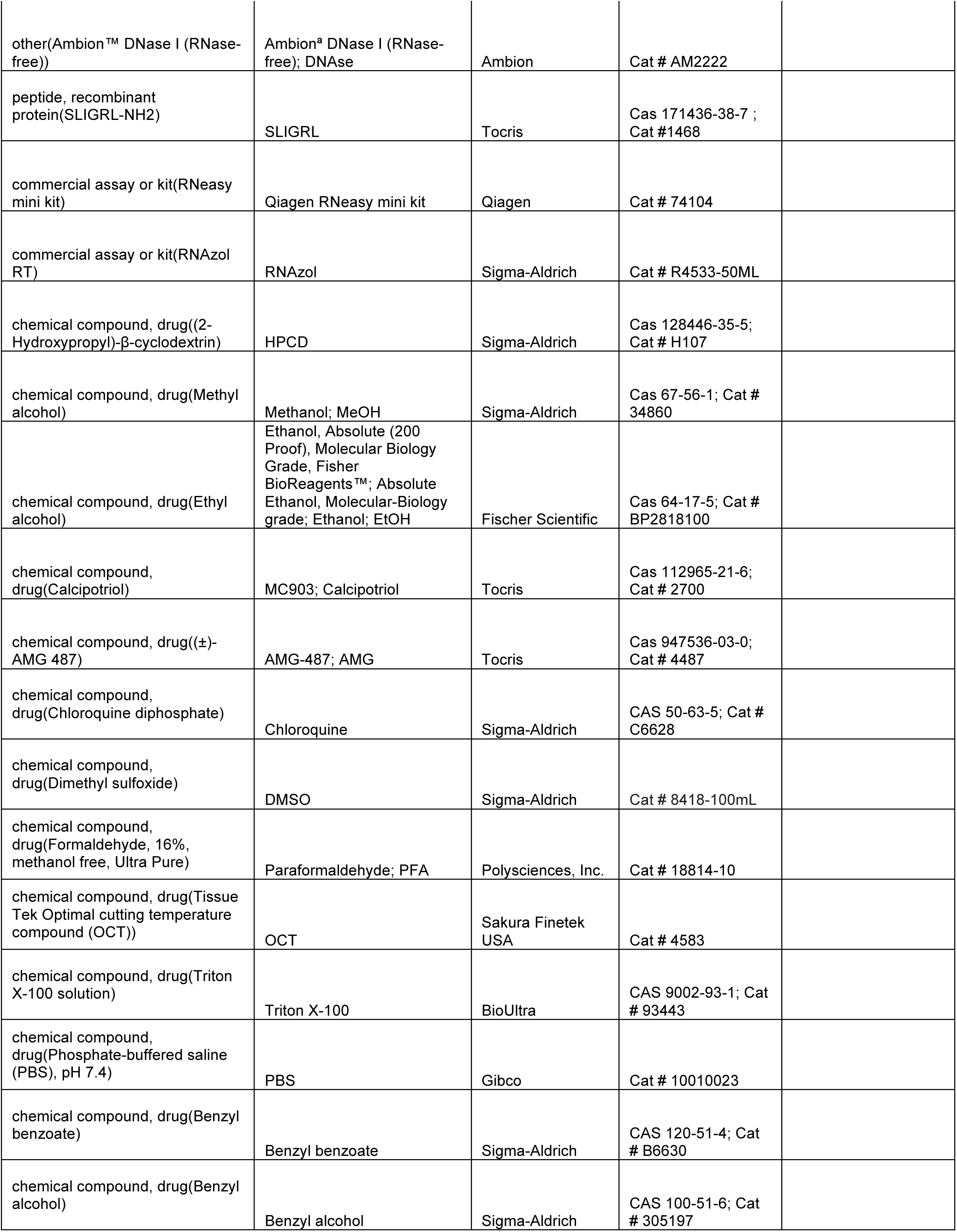
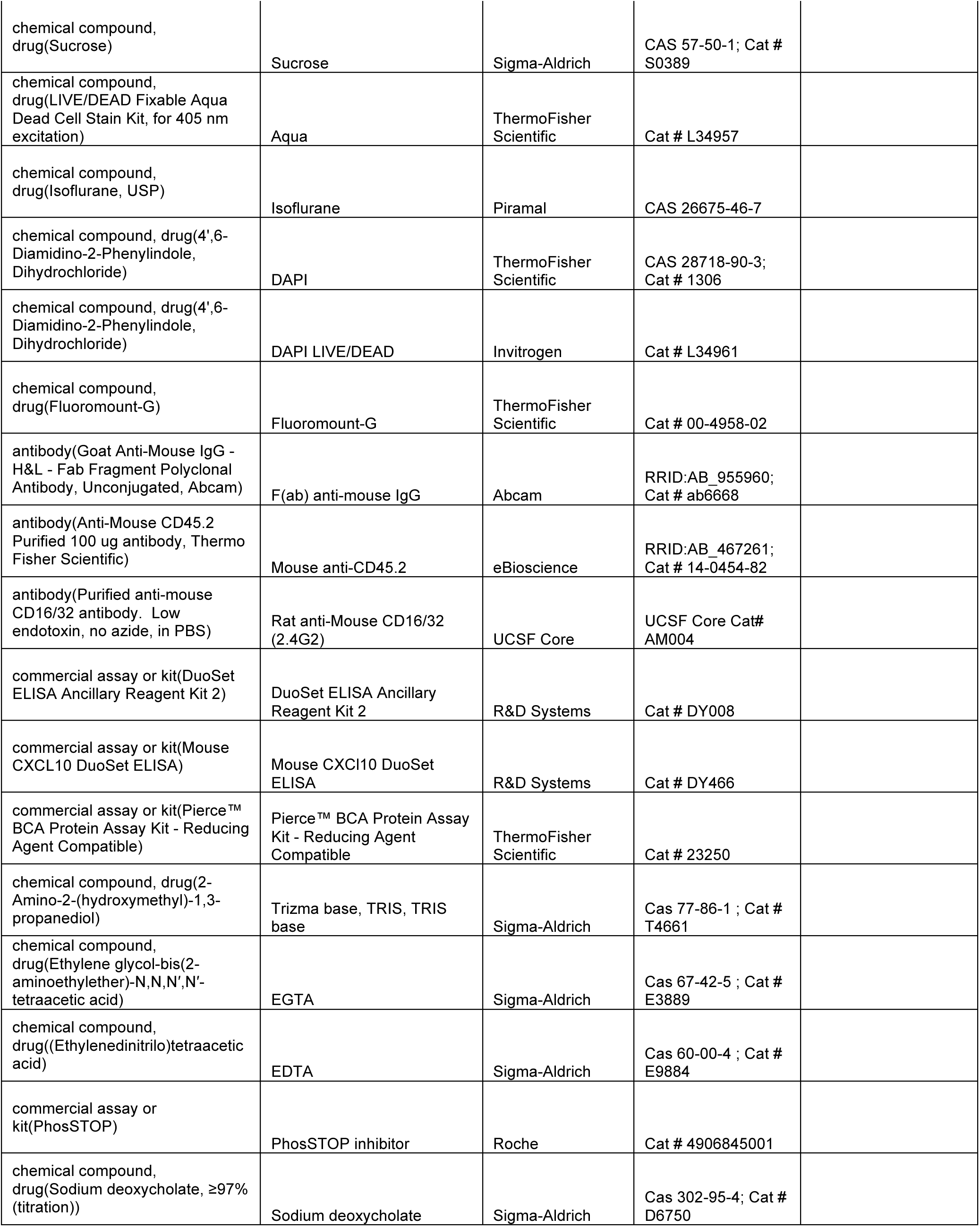
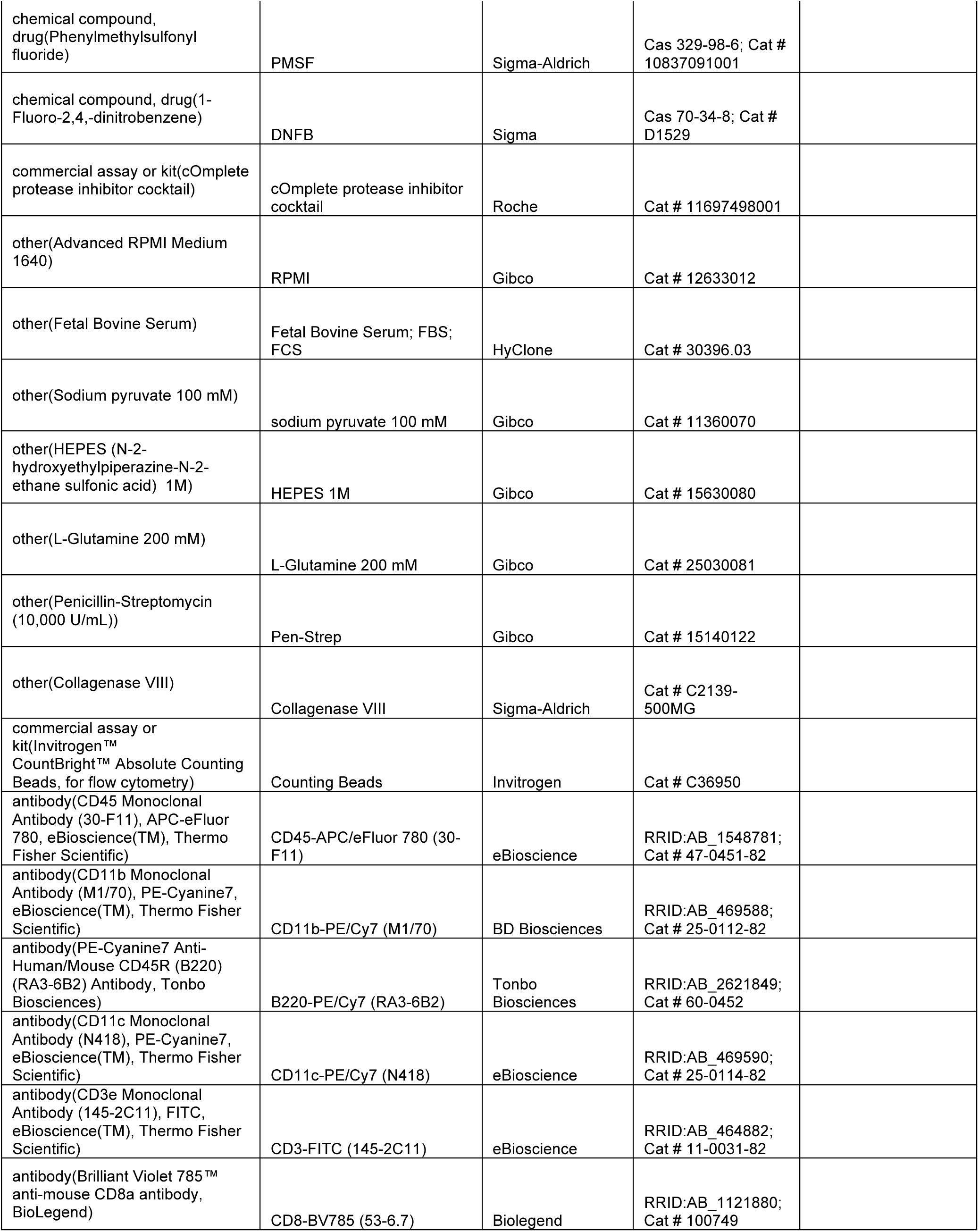
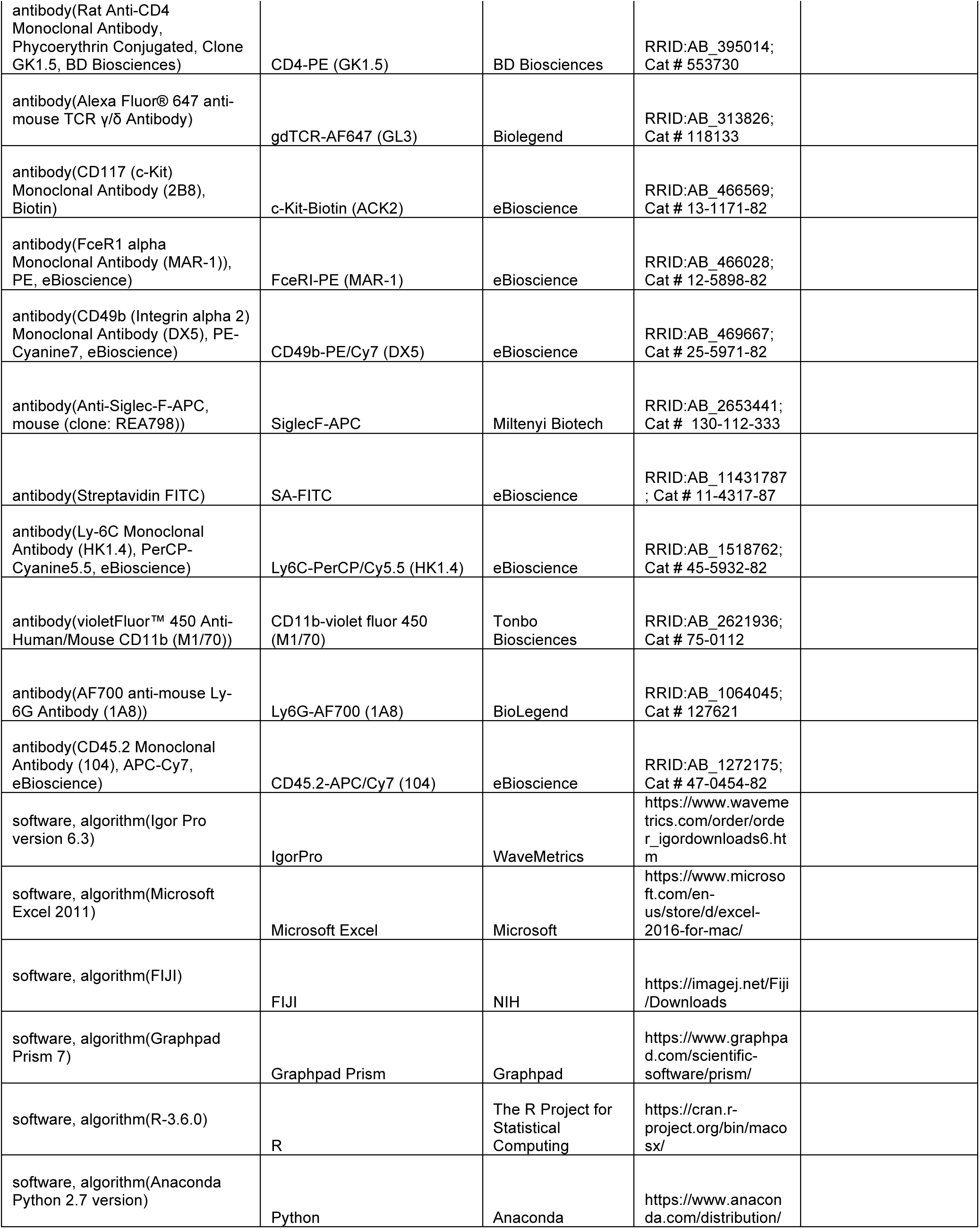
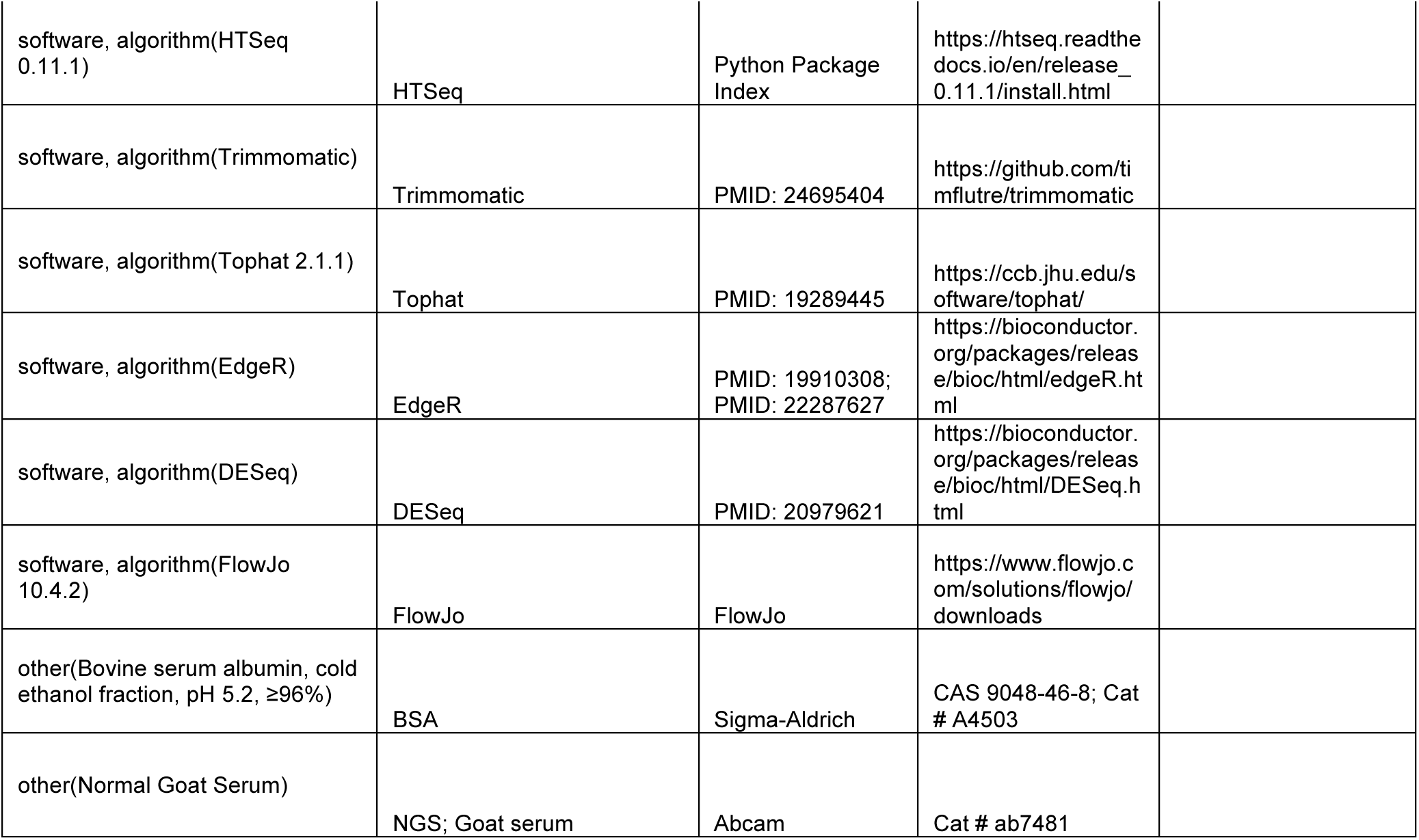

### Mouse studies

All mice were housed in standard conditions in accordance with standards approved by the Animal Care and Use Committee of the University of California Berkeley (12 hr light-dark cycle, 21°C). Wild-type C57BL/6 mice were obtained from Charles River or Jackson Laboratories and raised in-house. TSLPR^-/-^ mice were kindly provided by Dr. Steven Ziegler (*Crlf2^tm1Jni^*^59^)and backcrossed onto C57BL/6. All experiments were performed under the policies and recommendations of the International Association for the Study of Pain and approved by the University of California Berkeley Animal Care and Use Committee. Where appropriate, genotypes were assessed using standard PCR.

### MC903 model of atopic dermatitis

MC903 (Calcipotriol; R&D Systems) was applied to the shaved mouse cheek (20 µl of 0.2 mM in ethanol) or rostral back (40 µl of 0.2 mM in ethanol) once per day for 1-12 days using a pipette. 100% ethanol was used. All MC903 studies were performed on 8-12 week old age-matched mice. Behavior, RNA-seq, flow cytometry, and immunohistochemistry were performed on days 1, 2, 3, 5, 8 and/or 12. For AMG 487 experiments in the MC903 model, 50 µL 3.31 mM AMG 487 (Tocris) or 20% HPCD-PBS vehicle was injected subcutaneously one hour prior to recording behavior.^75^ Spontaneous scratching was manually scored for the first 30 minutes of observation. Both bout number and length were recorded. Behavioral scoring was performed while blind to experimental condition and mouse genotype.

### MC903 RNA isolation and sequencing

On days 1 (six hours post-treatment), 2, 5, or 8 post-treatment, mice treated with MC903 and vehicle were euthanized via isoflurane and cervical dislocation. Cheek skin was removed, flash-frozen in liquid nitrogen, and cryo-homogenized with a mortar and pestle. Ipsilateral trigeminal ganglia were dissected and both skin and trigeminal ganglia were homogenized for three minutes (skin) or one minute (TG) in 1 mL RNAzol RT (Sigma-Aldrich). Thoracic spinal cord was dissected from mice treated with 40 µL MC903 or ethanol on the shaved rostral back skin and homogenized for one minute in 1 mL RNAzol. Large RNA was extracted using RNAzol RT per manufacturer’s instructions. RNA pellets were DNase treated (Ambion), resuspended in 50 µL DEPC-treated water, and subjected to poly(A) selection and RNA-seq library preparation (Apollo 324) at the Functional Genomics Laboratory (UC Berkeley). Single-end read sequencing (length = 50 bp) was performed by the QB3 Vincent G. Coates Genomic Sequencing Laboratory (UC Berkeley) on an Illumina HiSeq4000. See **Supplementary Table 1** for number of mice per experimental condition and number of mapped reads per sample. Data are available at Gene Expression Omnibus under GSE132173.

### MC903 RNA sequencing analysis

Reads were mapped to the mm10 mouse genome using Bowtie2 and Tophat, and reads were assigned to transcripts using htseq-count.^113, 114^ For a given time point, replicate measurements for each gene from treated and control mice were used as input for DESeq (R) and genes with *p*_adjusted_ < 0.05 (for skin and spinal cord) or *p*_adjusted_ < 0.1 (for trigeminal ganglia) for at least one time point were retained for analysis.^115, 116^ For the skin dataset, we collated a set of AD-related immune cell markers, cytokines, atopic dermatitis disease genes, neurite outgrown/axonal guidance genes, and locally expressed neuronal transcripts, and from this list visualized genes that were significantly differentially expressed for at least one time point. For the trigeminal ganglia dataset, we plotted all genes that were significantly differentially expressed for at least one time point. Genes from these lists were plotted with hierarchical clustering using heatmap2 (R).

### Custom gene groups

Genes were clustered into functional groups and significance was evaluated using a permutation test. Briefly, we first tabulated the absolute value of the log_2_ fold change of gene expression (between MC903 and EtOH) of each gene in a given group of *n* genes in turn, and then we calculated the median of these fold change values, *z_true_*. We then drew *n* random genes from the set of all genes detected in the samples and computed the median log_2_ fold change as above using this null set, *z_null_*. Repeating the latter 10,000 times established a null distribution of median log_2_ fold change values; we took the proportion of resampled gene groups that exhibited (*z_true_* ≥ *z_null_*) as an empirical *p*-value reporting the significance of changes in gene expression for a given group of *n* genes.

### Flow Cytometry

Skin samples were collected from the cheek of mice at the indicated time points with a 4- or 6-mm biopsy punch into cold RPMI 1640 medium (RPMI; Gibco) and minced into smaller pieces with surgical scissors. When ear skin was collected, whole ears were dissected postmortem into cold RPMI and finely minced with scissors. For isolation of immune cells, skin samples were digested for 1h at 37°C using 1 U/mL Liberase TM (Roche) and 5 µg/mL DNAse I (Sigma). At the end of the digestion, samples were washed in FACS buffer (PBS with 0.5% FCS and 2 mM EDTA) and filtered through a 70 or 100 µm strainer (Falcon). Cells were stained with LIVE/DEAD fixable stain Aqua in PBS (Invitrogen), then blocked with anti-CD16/32 (UCSF Core) and stained with the following fluorophore-conjugated antibodies (all from eBiosciences unless stated otherwise) in FACS buffer: cKit-Biotin (clone ACK2; secondary stain with SA-FITC), CD11b-violet fluor 450 (Tonbo; clone M1/70), Ly6C-PerCP/Cy5.5 (clone HK1.4), CD49b-PE/Cy7 (clone DX5), CD45.2-APC/Cy7 (clone 104), FceRI-PE (MAR-1), Ly6G-AF700 (clone 1A8). 10 µL of counting beads (Invitrogen) were added after the last wash to measure absolute cell counts. For measurement of CD4^+^ T cells, 6-mm skin biopsy punch samples were digested for 30 minutes at 37°C using Collagenase VIII (Sigma). At the end of the digestion, cells were washed in RPMI buffer (RPMI with: 5% FCS, 1% penicillin-streptomycin, 2 mM L-glutamine, 10 mM HEPES buffer, 1 mM sodium pyruvate). Cells were blocked with anti-CD16/32 (UCSF Core) and stained with the following fluorophore-conjugated antibodies in FACS buffer (PBS with 5% FCS and 2 mM EDTA): CD45-APC-eFluor780 (clone 30-F11; eBiosciences), CD11b-PE/Cy7 (clone M1/70; BD Biosciences), B220-PE/Cy7 (clone RA3-6B2; Tonbo Biosciences), CD11c-PE/Cy7 (clone N418; eBiosciences), CD3-FITC (clone 145-2C11; eBiosciences), CD8-BV785 (clone 53-6.7; Biolegend), CD4-PE (clone GK1.5; BD Biosciences), gdTCR-AF647 (clone GL3; Biolegend). 10 µL of counting beads (Invitrogen) were added after the last wash to measure absolute cell counts, and samples were resuspended in DAPI LIVE/DEAD (Invitrogen). Blood samples were collected from saphenous vein or from terminal bleed following decapitation. Red blood cells were lysed using ACK lysis buffer (Gibco), and samples were washed with FACS buffer (PBS with 0.5% FCS and 2 mM EDTA), and blocked with anti-CD16/32. Cells were stained with Ly6G-PE (1A8; BD Biosciences), CD11b-violet fluor 450 (M1/70, Tonbo), Ly6C-PerCP/Cy5.5 (HK1.4, Biolegend), and aGr1-APC/Cy7 (RB6-8C5, eBiosciences). For all experiments, single cell suspensions were analyzed on an LSR II or LSR Fortessa (BD Biosciences), and data were analyzed using FlowJo (TreeStar, v.9.9.3) software.

### Human keratinocyte RNA sequencing

Normal human epidermal keratinocytes from juvenile skin (PromoCell #C-12001) were cultured in PromoCell Keratinocyte Growth Medium 2 and passaged fewer than 5 times. Cells were treated for three hours at room temperature with 100 µM SLIGRL or vehicle (Ringer’s + 0.1% DMSO). Total RNA was extracted by column purification (Qiagen RNeasy Mini Kit). RNA was sent to the Vincent J. Coates Sequencing Laboratory at UC Berkeley for standard library preparation and sequenced on an Illumina HiSeq2500 or 4000. Sequences were trimmed (Trimmomatic), mapped (hg19, TopHat) and assigned to transcripts using htseq-count. Differential gene expression was assessed using R (edgeR). Data are available at Gene Expression Omnibus under GSE132174.

### IHC of whole-mount skin

Staining was performed as previously described.^117, 118^ Briefly, 8-week old mice were euthanized and the cheek skin was shaved. The removed skin was fixed overnight in 4% PFA, then washed in PBS (3X for 10 min each). Dermal fat was scraped away with a scalpel and skin was washed in PBST (0.3% Triton X-100; 3X for two hours each) then incubated in 1:500 primary antibody (Rabbit anti beta-Tubulin II; Abcam #ab18207 or Rabbit anti-CGRP; Immunostar #24112) in blocking buffer (PBST with 5% goat serum and 20% DMSO) for 6 days at 4°C. Skin was washed as before and incubated in 1:500 secondary antibody (Goat anti-Rabbit Alexa 594; Invitrogen #R37117) in blocking buffer for 3 days at 4°C. Skin was washed in PBST, serially dried in methanol: PBS solutions, incubated overnight in 100% methanol, and finally cleared with a 1:2 solution of benzyl alcohol: benzyl benzoate (BABB; Sigma) before mounting between No. 1.5 coverglass. Whole mount skin samples were imaged on a Zeiss LSM 880 confocal microscope with OPO using a 20x water objective. Image analysis was performed using a custom macro in FIJI. Briefly, maximum intensity z-projections of the beta-tubulin III or CGRP channel were converted to binary files that underwent edge-detection analysis. Regions were defined by circling all stained regions. Region sizes and locations were saved.

### IHC of sectioned trigeminal ganglia

TG were dissected from 8- to 12-week old adult mice and post-fixed in 4% PFA for one hour. TG were cryo-protected overnight at 4°C in 30% sucrose-PBS, embedded in OCT, and then cryosectioned at 14 µm onto slides for staining. Slides were washed 3x in PBST (0.3% Triton X-100), blocked in 2.5% Normal Goat serum + 2.5% BSA-PBST, washed 3X in PBST, blocked in endogenous IgG block (1:10 F(ab) anti-mouse IgG (Abcam ab6668) + 1:1000 Rat anti-mouse CD16/CD32 (UCSF MAB Core) in 0.3% PBST), washed 3X in PBST and incubated overnight at 4°C in 1:1000 primary antibody in PBST + 0.5% Normal Goat Serum + 0.5% BSA. Slides were washed 3x in PBS, incubated 2 hr at RT in 1:1000 secondary antibody, washed 3X in PBS, and then incubated 30 min in 1:2000 DAPI-PBS. Slides were washed 3x in PBS and mounted in Fluoromount-G with No. 1.5 coverglass. Primary antibodies used: Mouse anti-CD45 (eBioscience #14-054-82) and Chicken anti-Peripherin (Abcam #39374). Secondary antibodies used: Goat anti-Chicken Alexa 594 (ThermoFisher #A11042) and Goat anti-Mouse Alexa 488 (Abcam #150117). DAPI (ThermoFisher #D1306) was also used to mark nuclei. Imaging of TG IHC experiments was performed on an Olympus IX71 microscope with a Lambda LS-xl light source (Sutter Instruments). For TG IHC analysis, images were analyzed using automated scripts in FIJI (ImageJ) software. Briefly, images were separated into the DAPI, CD45, and Peripherin channels. The minimum/maximum intensity thresholds were batch-adjusted to pre-determined levels, and adjusted images were converted to binary files. Regions were defined by circling all stained regions with pre-determined size-criteria. Region sizes and locations were saved. All scripts are available upon request.

### Neutrophil depletion

Neutrophils were acutely depleted using intraperitoneal injection with 250 µg aGR1 in PBS (clone RB6-8C5, a gift from D. Portnoy, UC Berkeley, or from Biolegend), 16-24 hours before behavioral and flow cytometry experiments. Depletion was verified using flow cytometry on blood collected from terminal bleed following decapitation. For longer depletion experiments using the MC903 model, mice were injected (with 250 µg aGR1 in PBS or PBS vehicle, i.p.) beginning one day prior to MC903 administration and each afternoon thereafter through day 7 of the model, or on days 8-11 for measurement of day 12 itch behaviors, and blood was collected via saphenous venipuncture at days 3, 5, or by decapitation at day 8 to verify depletion.

### CXCL10 ELISA measurements in skin

Neutrophil-depleted or uninjected mice were treated with MC903 or ethanol for 7 days. On day 8, 6-mm biopsy punches of cheek skin were harvested, flash-frozen in liquid nitrogen, cryo-homogenized by mortar and pestle, and homogenized on ice for three minutes at maximum speed in 0.5 mL of the following tissue homogenization buffer (all reagents from Sigma unless stated otherwise): 100 mM Tris, pH 7.4; 150 mM NaCl, 1 mM EGTA, 1 mM EDTA, 1% Triton X-100, and 0.5% Sodium deoxycholate in ddH2O; on the day of the experiment, 200 mM fresh PMSF in 100% ethanol was added to 1mM, with 1 tablet cOmplete protease inhibitor (Roche) per 50 mL, and 5 tablets PhosSTOP inhibitor (Roche) per 50 mL buffer. Tissues were agitated in buffer for two hours at 4°C, and centrifuged at 13,000 rpm for 20 minutes at 4°C. Supernatants were aliquoted and stored at −80°C for up to one week. After thawing, samples were centrifuged at 10,000 rpm for five minutes at 4°C. Protein content of skin homogenates was quantified by BCA (Thermo Scientific) and homogenates were diluted to 2 mg/mL protein in PBS and were subsequently diluted 1:2 in Reagent Diluent (R&D Systems). CXCL10 protein was quantified using the Mouse CXCL10 Duoset ELISA kit (R&D Systems; #DY466-05) according to manufacturer’s instructions. Plate was read at 450 nm and CXCL10 was quantified using a seven-point standard curve (with blank and buffer controls) and fitted with a 4-parameter logistic curve.

### Acute itch behavior

Itch behavioral measurements were performed as previously described.^56, 119, 120^ Mice were shaved one week prior to itch behavior and acclimated in behavior chambers once for thirty minutes at the same time of day on the day prior to the experiment. Behavioral experiments were performed during the day. Compounds injected: 1 µg carrier-free CXCL1 (R&D systems) in PBS, 3.31 mM AMG 487 (Tocris, prepared from 100 mM DMSO stock) in 20% HPCD-PBS, 50 mM Chloroquine diphosphate (Sigma) in PBS, along with corresponding vehicle controls. Acute pruritogens were injected using the cheek model (20 µL, subcutaneous/s.c.) of itch, as previously described.^56^ AMG 487 (50 µL) or vehicle was injected s.c. into the rostral back skin one hour prior to recording of behavior. Behavioral scoring was performed as described above.

### Lipidomics

Skin was collected from the cheek of mice post-mortem with a 6-mm biopsy punch and immediately flash-frozen in liquid nitrogen. Lipid mediators and metabolites were quantified via liquid chromatography-tandem mass spectrometry (LC-MS/MS) as described before.^121^ In brief, skin was homogenized in cold methanol to stabilize lipid mediators. Deuterated internal standards (PGE_2_-d4, LTB_4_-d4, 15-HETE-d8, LXA_4_-d5, DHA-d5, AA-d8) were added to samples to calculate extraction recovery. LC-MS/MS system consisted of an Agilent 1200 Series HPLC, Luna C18 column (Phenomenex, Torrance, CA, USA), and AB Sciex QTRAP 4500 mass spectrometer. Analysis was carried out in negative ion mode, and lipid 30 mediators quantified using scheduled multiple reaction monitoring (MRM) mode using four to six specific transition ions per analyte.^122^

### 1-Fluoro-2,4-dinitrobenzene (DNFB) model of atopic dermatitis

The DNFB model was conducted as described previously.^63^ Briefly, the rostral backs of isofluorane-anesthetized mice were shaved using surgical clippers. Two days after shaving, mice were treated with 25 µL 0.5% DNFB (Sigma) dissolved in 4:1 acetone:olive oil vehicle on the rostral back using a pipette. Five days after the initial DNFB sensitization, mice were challenged with 40 µL 0.2% DNFB or 4:1 acetone:olive oil vehicle applied to the outer surface of the right ear. Twenty-four hours after DNFB or vehicle challenge, mice were euthanized and ear skin was harvested for flow cytometry.

### Statistical analyses

Different control experimental conditions (*e.g.* uninjected versus PBS-injected animals) were pooled when the appropriate statistical test showed they were not significantly different (Supplementary Table 2). For all experiments except RNA-seq (see above), the following statistical tests were used, where appropriate: Student’s t-test, one-way ANOVA with Tukey-Kramer post hoc comparison, and two-way ANOVA with Tukey Kramer or Sidak’s post-hoc comparison. Bar graphs show mean ± SEM. Statistical analyses were performed using PRISM 7 software (GraphPad). For all *p* values, *=0.01<*p*<0.05, **=0.001<*p*<0.01, ***=0.0001<*p*<0.001, and ****=*p*<0.0001.

## Acknowledgements

The authors would like to thank members of the Bautista and Barton labs for helpful discussions on the data. We are grateful to S. Ziegler (Ben Aroya Research Institute) for the gift of the TSLPR^-/-^ mouse. We also thank M. Pellegrino and L. Thé for pilot studies on human keratinocyte transcriptome, and T. Morita and J. Wong for technical assistance with mouse behavioral experiments. D.M.B. is supported by the NIH (AR059385; NS07224 and NS098097, also to R.B.B) and the Howard Hughes Medical Institute. G.M.B. is supported by the NIH (AI072429, AI063302, AI104914, AI105184) and the Burroughs Wellcome Fund. J.D. was supported by a Long-Term Fellowship from the Human Frontier Science Program (LT-000081/2013-L). K.G. is supported by NIH grant EY026082. Confocal imaging experiments were conducted at the CRL Molecular Imaging Center, supported by the Helen Wills Neuroscience Institute (UC Berkeley). We would like to thank H. Aaron and F. Ives for their microscopy training and assistance. This work used the Functional Genomics Laboratory and the Vincent J. Coates Genomics Sequencing Laboratory at UC Berkeley, supported by NIH S10 OD018174 Instrumentation Grant.

---

**Figure 1-Figure Supplement 1.**
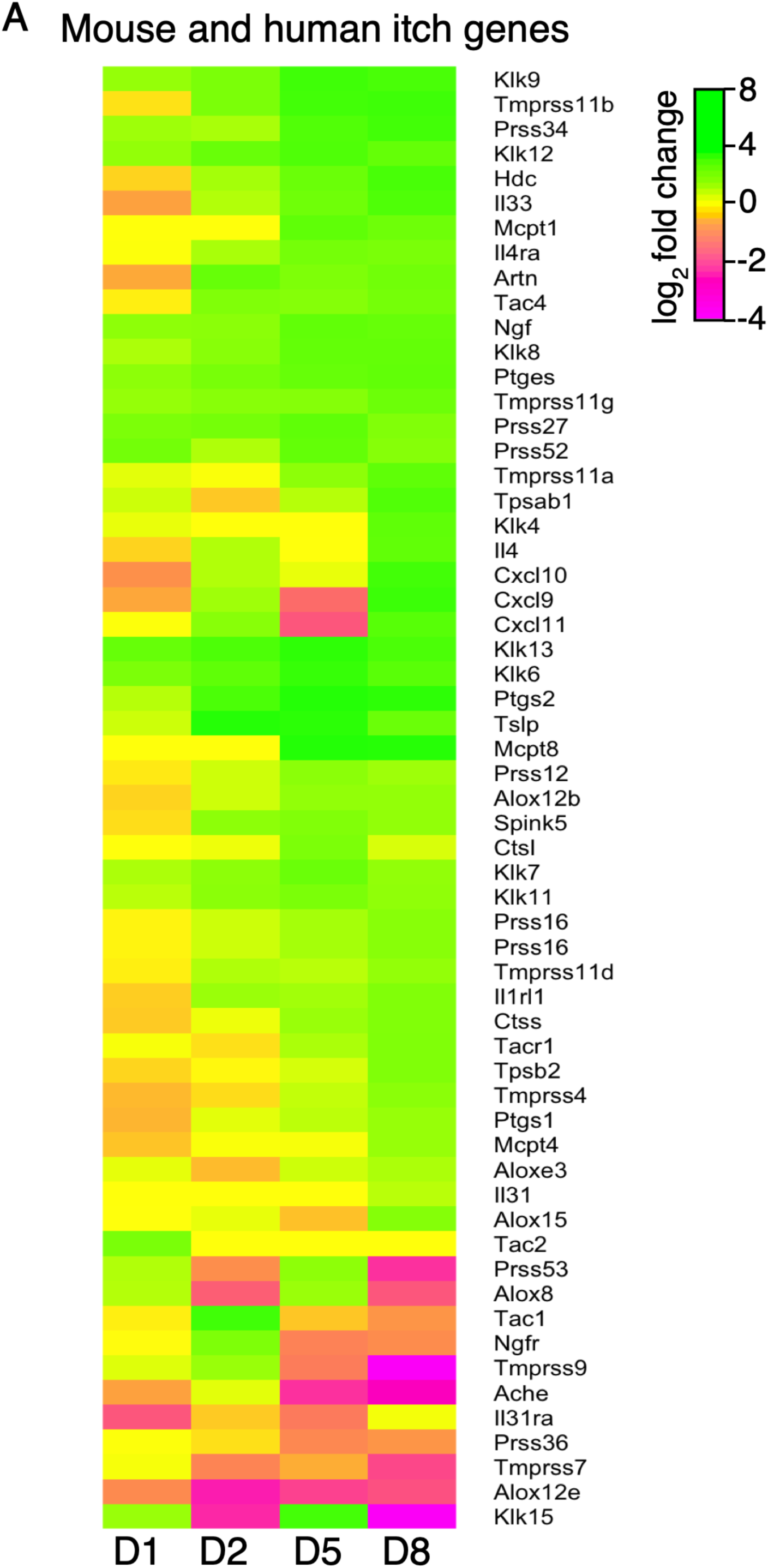
Expression of mouse and human itch genes. **A.** Log_2_ fold change in gene expression (MC903 vs. ethanol) in mouse skin at indicated time points for genes implicated in mouse or human acute or chronic itch that were significantly differentially expressed for at least one time point in the MC903 model. Green bars = increased expression in MC903 relative to ethanol; magenta = decreased expression. Exact values and corrected *p*-values are reported in **Figure 1-source data 6** and **Supplemental Data**, respectively.

**Figure 1-Figure Supplement 2.**
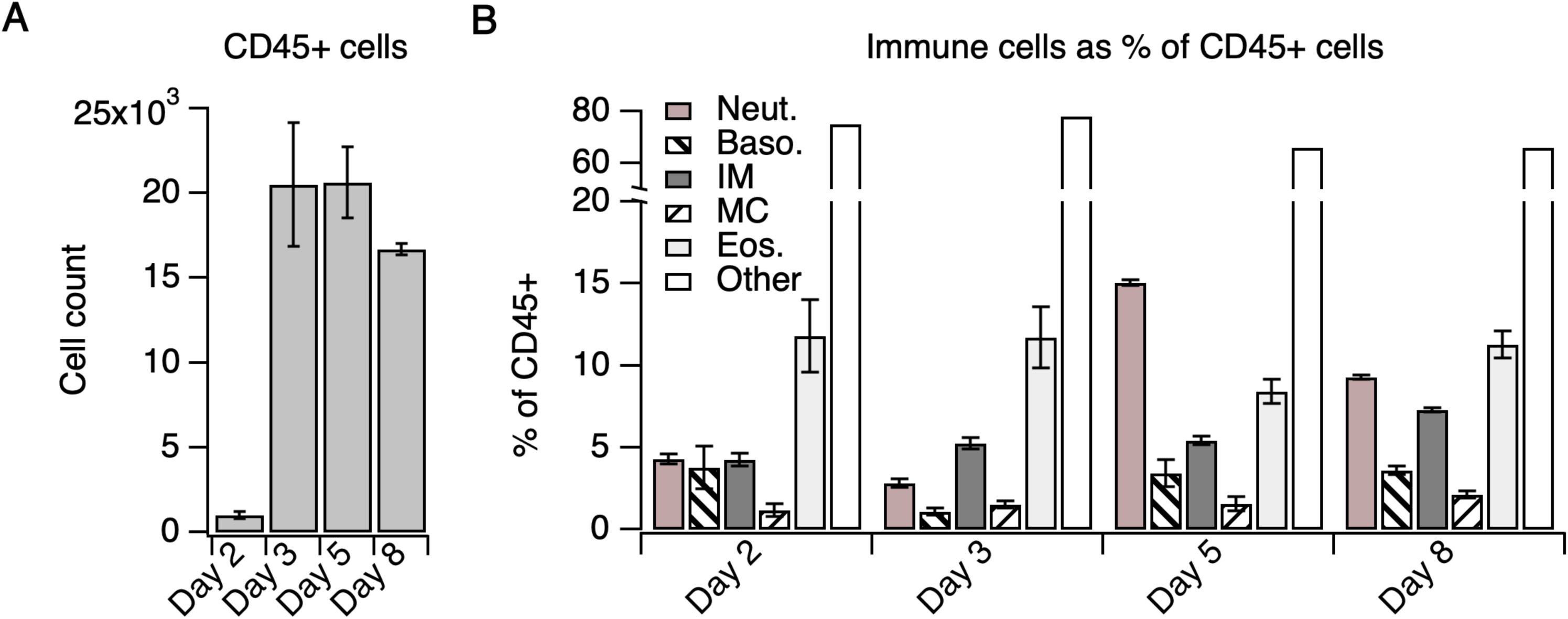
Immune cells represented as % of CD45^+^ cells. **A.** Number of CD45^+^ cells in MC903-treated skin on days 2-8 of the model. **B.** Skin-infiltrating immune cell subtypes on days 2-8 of the MC903 model shown in Figure 1, represented as % of CD45^+^ cells. CD4^+^ T cell measurements were acquired using a separate staining panel from different animals than the myeloid cell measurements (see Methods) and were not included. See **Supplementary Table 3** for % of CD45^+^ cell measurements for all flow cytometry experiments. Error bars represent mean ± SEM.

**Figure 1-Figure Supplement 3.**
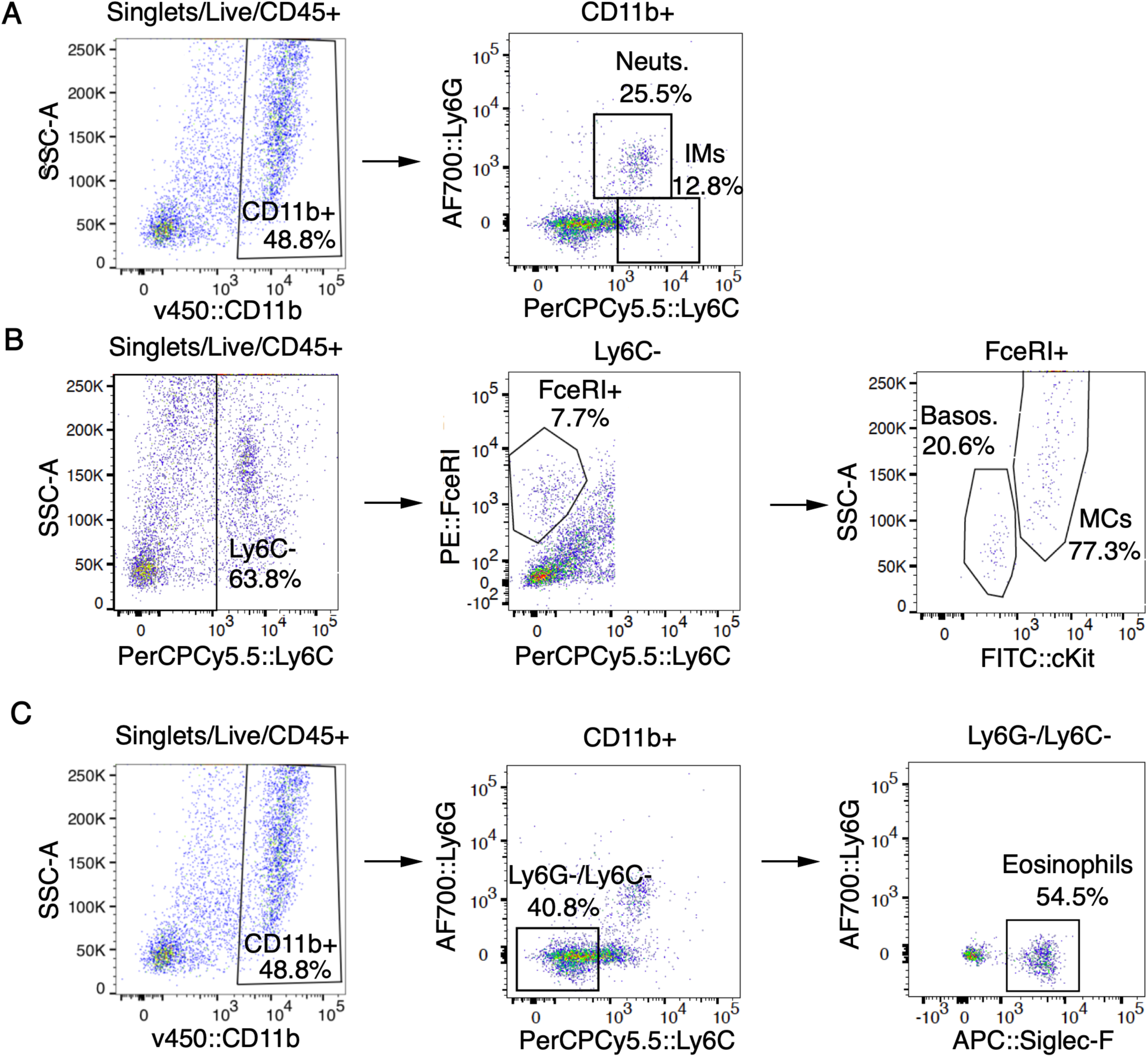
Myeloid and granulocyte gating strategy. **A-C.** Representative FACS plots of cells isolated from MC903-treated cheek skin showing gating strategy for neutrophils (A), inflammatory monocytes (A), mast cells (B), basophils (B), and eosinophils (C) as shown in Figure 1E-F and Figure 1-Figure Supplement 5.

**Figure 1-Figure Supplement 4.**
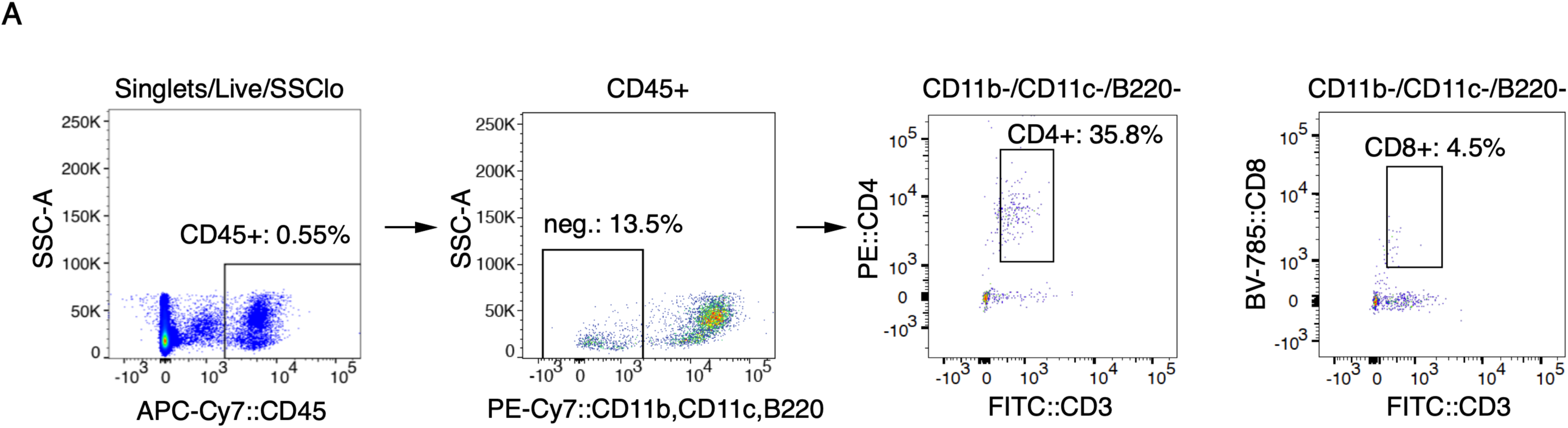
T cell gating strategy. **A.** Representative FACS plots of cells isolated from MC903-treated cheek skin showing gating strategy for CD4^+^ T cells as shown in Figure 1G.

**Figure 1-Figure Supplement 5.**
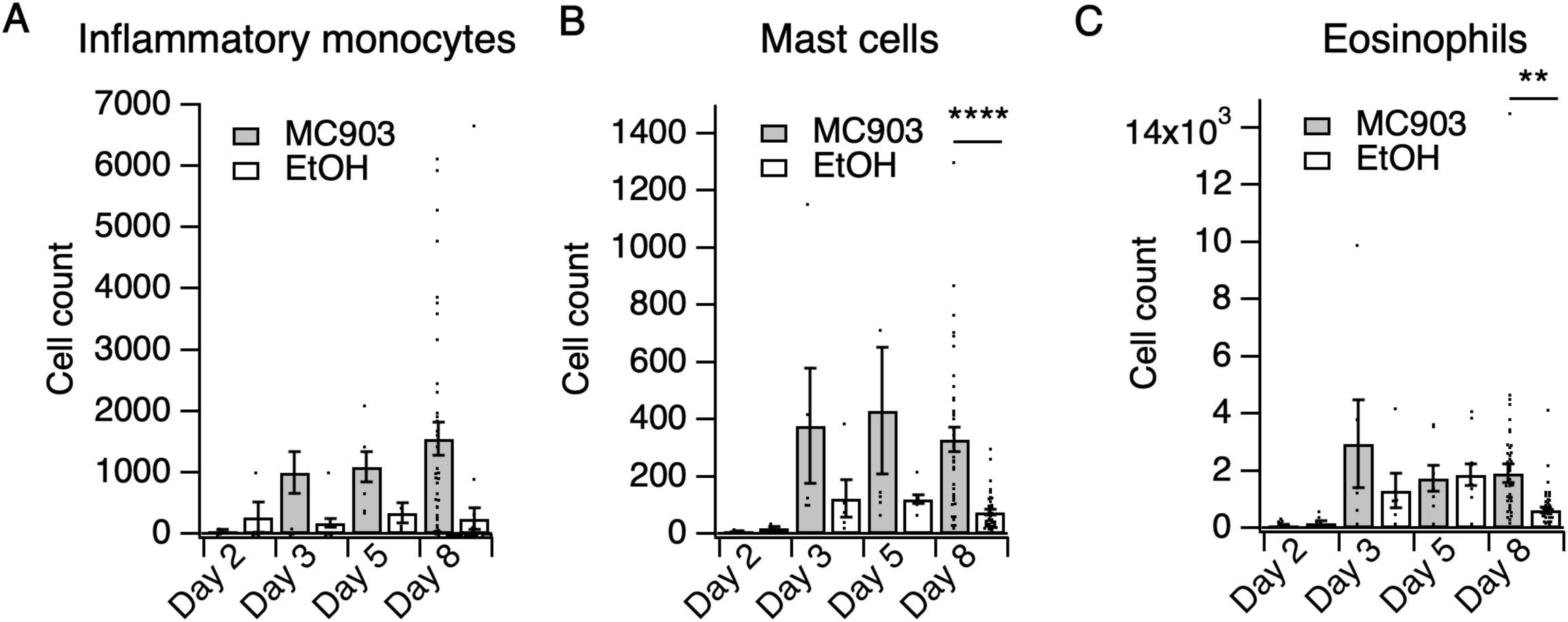
Immune cell counts in MC903-treated skin. **A.** Inflammatory monocyte counts in MC903- and ethanol-treated skin at indicated time points (two-way ANOVA: *p*_treatment_= 0.0662, F(1,102) = 3.44; n=4,4,5,5,6,8,40,38 mice). **B.** Mast cell counts in MC903- and ethanol-treated skin at indicated time points (two-way ANOVA: \*\**p*_treatment_= 0.0024, F(1,102) = 9.69; Sidak’s multiple comparisons: *p_day 2_* > 0.999, n=4,4 mice; *p_day 3_* = 0.3019, n=5,5 mice; *p_day 5_* = 0.0586, n=6,8 mice; \*\*\*\**p_day 8_* < 0.0001, n=40,38 mice). **C.** Eosinophil counts in MC903- and ethanol-treated skin at indicated time points (two-way ANOVA: *p*_time_= 0.0471, F(3,102) = 2.74; Sidak’s multiple comparisons: *p_day 2_* > 0.999, n=4,4 mice; *p_day 3_* = 0.3596, n=5,5 mice; *p_day 5_* = 0.9998, n=6,8 mice; \*\**p_day 8_* = 0.0020, n=40,38 mice). Data from mice receiving i.p. injection of PBS (see Figure 4) in addition to MC903 or EtOH are also included. Exact values displayed in **Figure 1-source data 4**.

**Figure 1-Figure Supplement 6.**
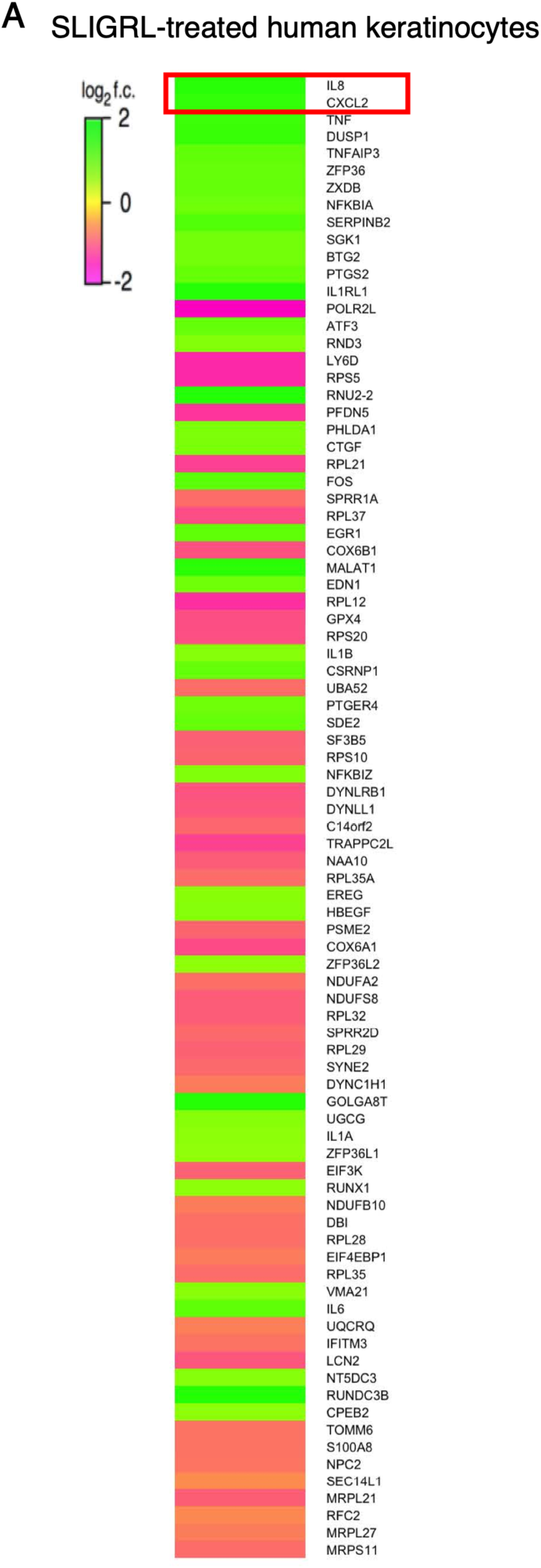
Protease receptor activation triggers rapid upregulation of neutrophil chemoattractant genes in human keratinocytes. **A.** Heat map showing log_2_ fold change in gene expression in cultured human keratinocytes 3 hours after SLIGRL treatment (100 µM; bottom; see **Figure 1-source data 7**) compared to vehicle controls, as measured by RNA-seq. Genes are sorted by descending corrected *p*-value; only significantly differentially expressed (*p* < 0.05) are displayed. Exact values and corrected *p*-values are reported in **Figure 1-source data 7** and **Supplemental Data**, respectively.

**Figure 1-Figure Supplement 7.**
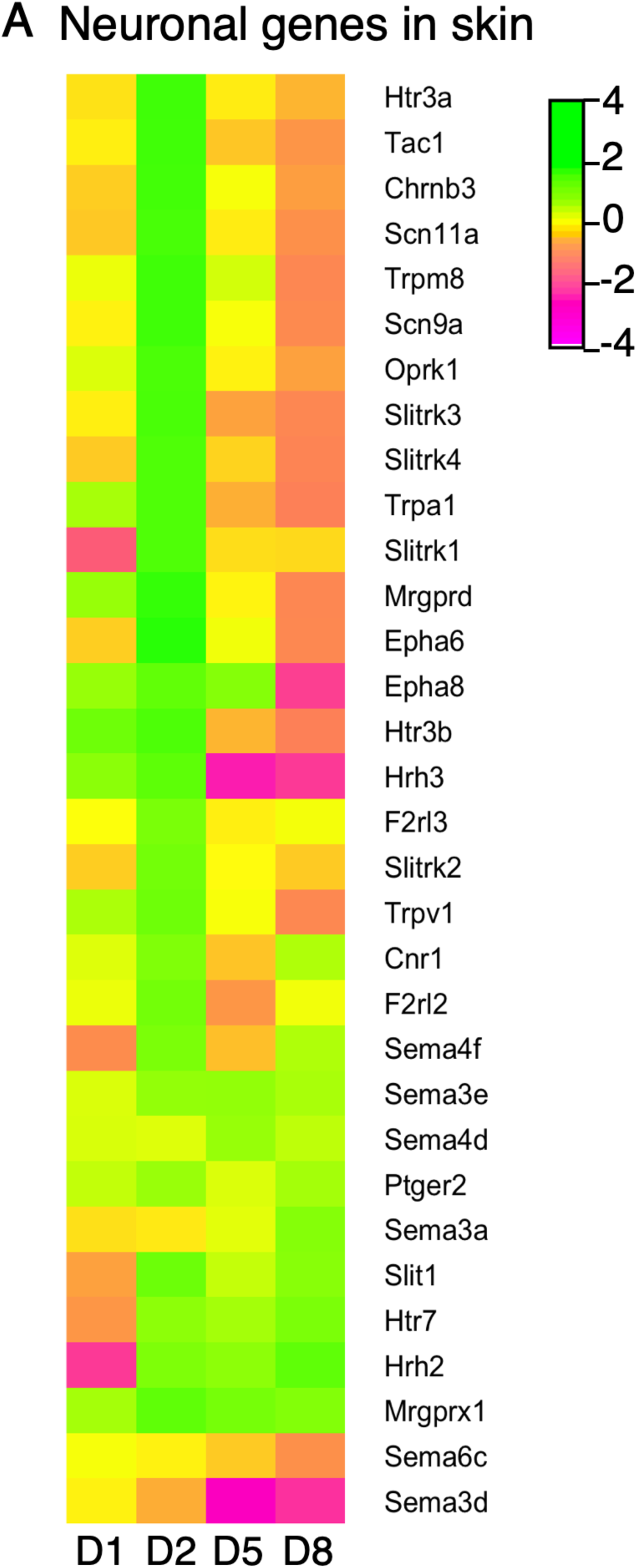
Expression of neuronal genes and axon guidance molecules in skin. **A.** Log_2_ fold change in gene expression (MC903 vs. EtOH) in mouse skin at indicated time points for markers of locally translated sensory neuronal transcripts or genes implicated in neurite remodeling and/or axon guidance that were significantly differentially expressed for at least one time point in the MC903 model. Green bars = increased expression in MC903 relative to ethanol; magenta = decreased expression. Exact values and corrected *p*-values are reported in **Figure 1-source data 8** and **Supplemental Data**, respectively.

**Figure 1-Figure Supplement 8.**
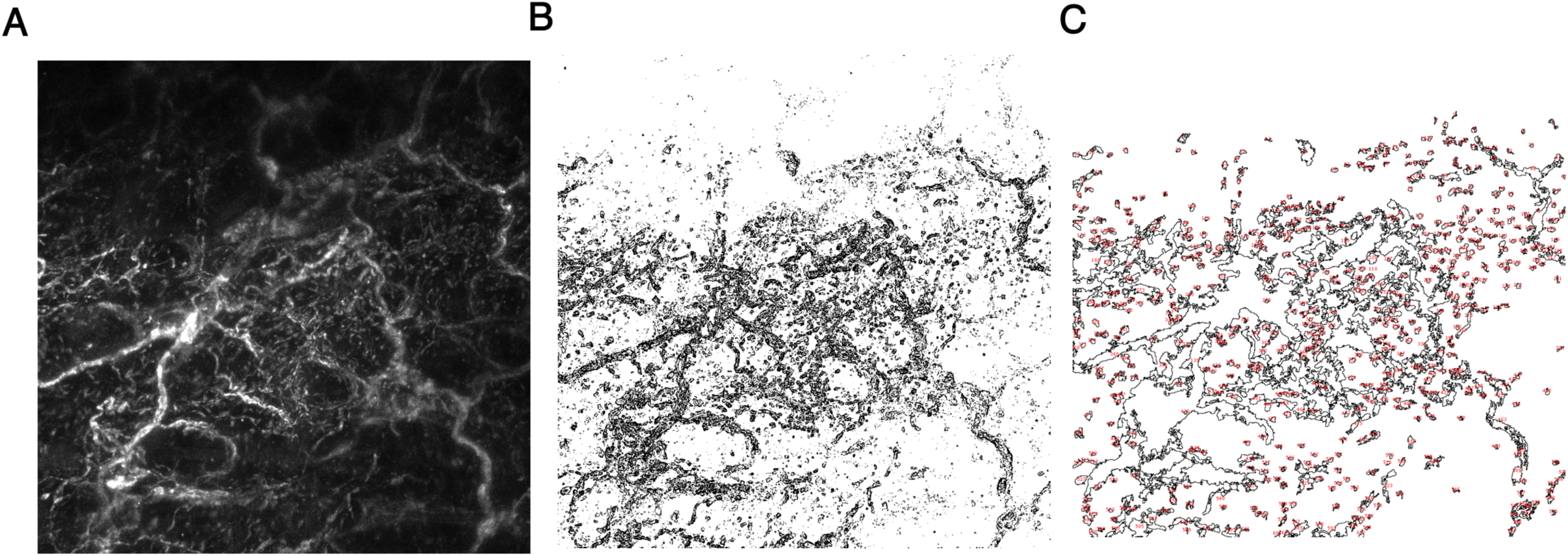
Method of image quantification for whole mount skin. **A.** Representative maximum intensity z-projection of beta tubulin III staining in cheek skin. **B.** Binary image after edge-detection. **C.** % Area innervated was calculated from the percentage of the image area which was occupied by the regions of interest (ROIs) outlined in red.

**Figure 1-Figure Supplement 9.**
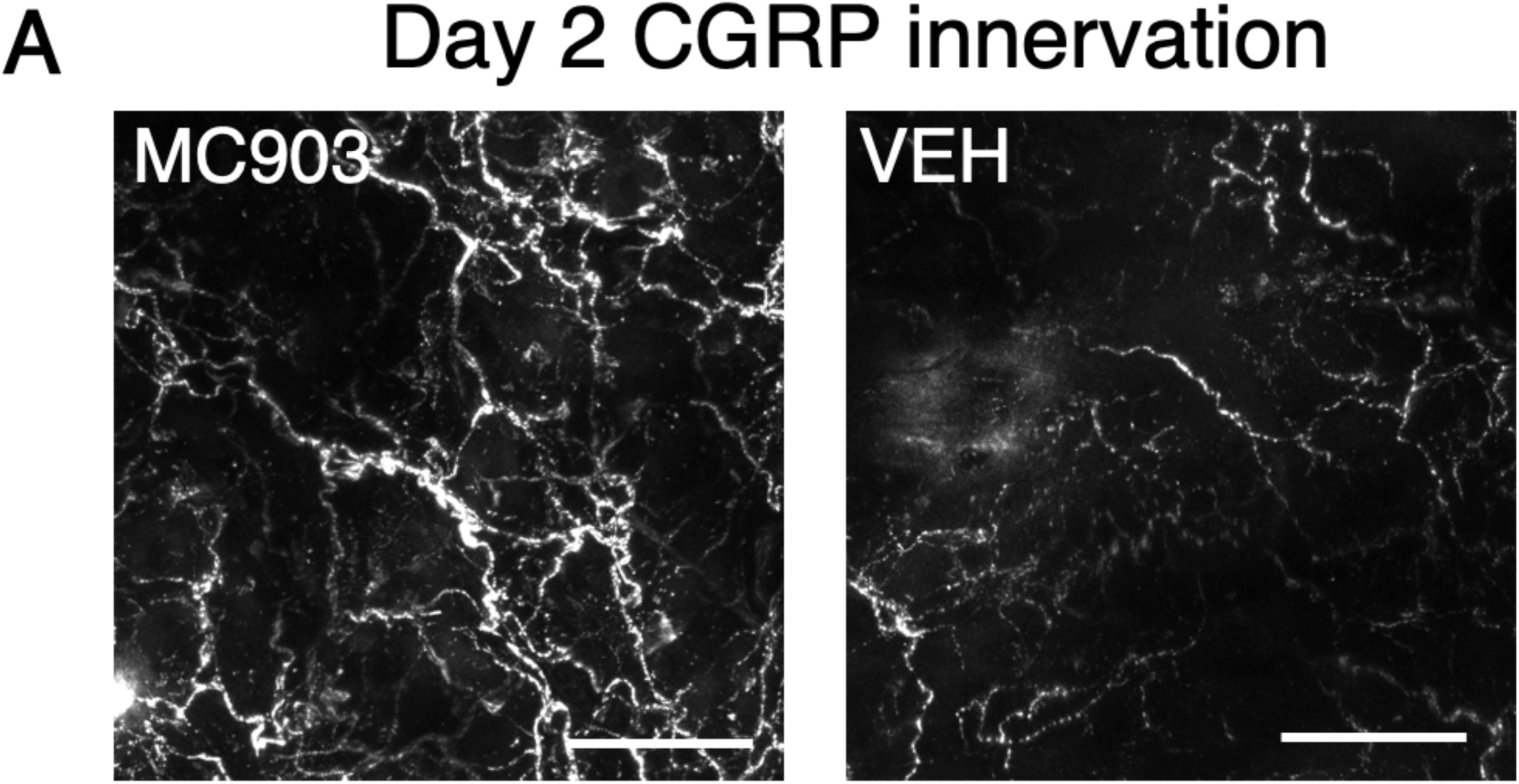
Peptidergic fibers display hyperinnervation in MC903-treated skin. **A.** Representative maximum intensity Z-projections from immunohistochemistry (IHC) of whole-mount mouse skin on day 2 of the MC903 model. Skin was stained with peptidergic neuronal marker Calcitonin related-gene peptide (CGRP; white). Images were acquired on a confocal microscope using a 20x water objective.

**Figure 1-Figure Supplement 10.**
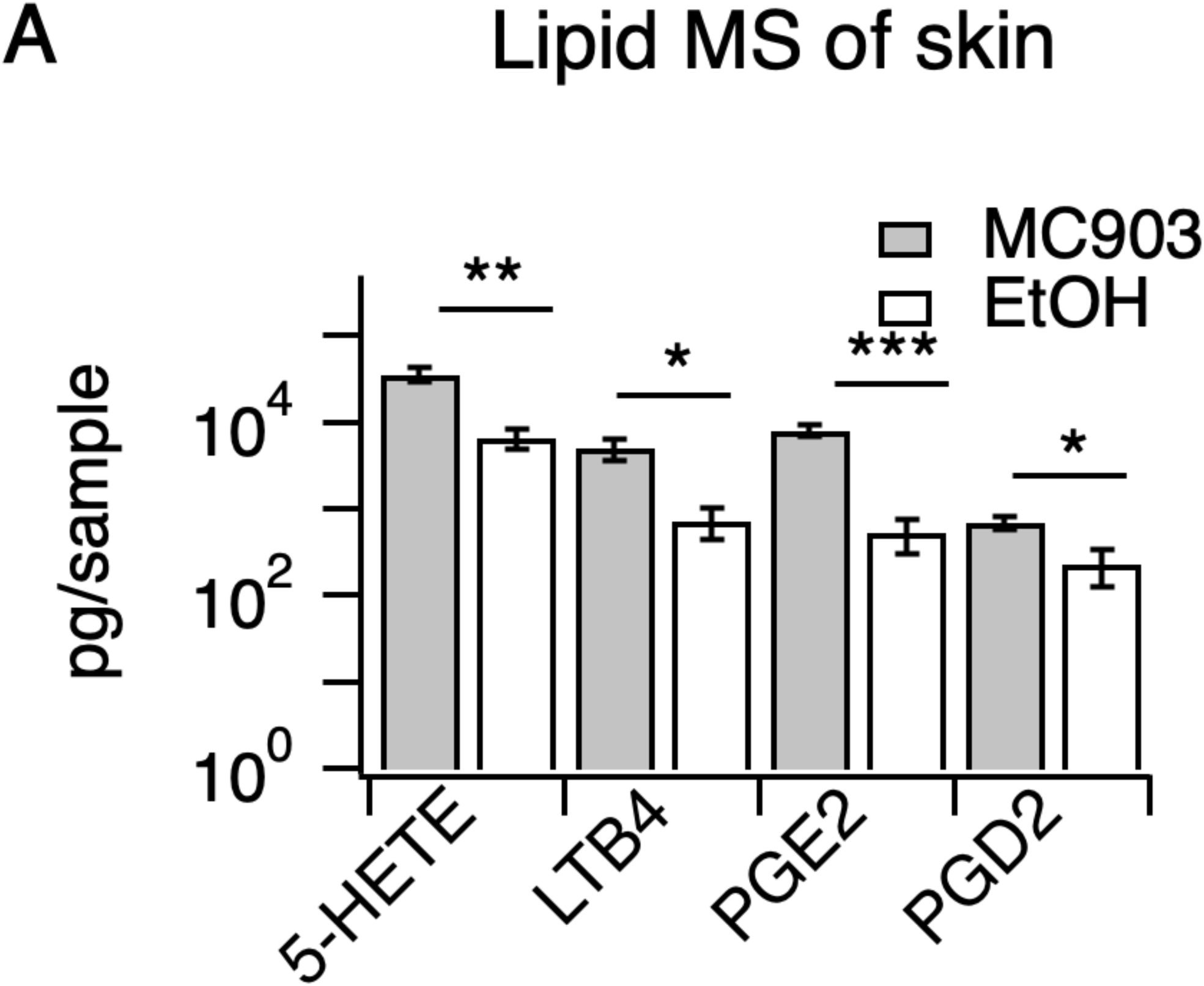
Inflammatory lipids in MC903-treated skin. **A.** Quantification of indicated lipids from 6 mm biopsy punches of cheek skin of MC903- and EtOH-treated mice (at day 8) by LC-MS/MS (**p = 0.006 (*t*=4.148,*df*=6), *p = 0.024 (*t*=3.003,*df*=6), ***p = 0.0007 (*t*=6.392,*df*=6), *p = 0.022 (*t*=3.058,*df*=6); two-tailed unpaired t-tests; n = 4 mice per group, see **Figure 1-source data 8**).

**Figure 2-Figure Supplement 1.**
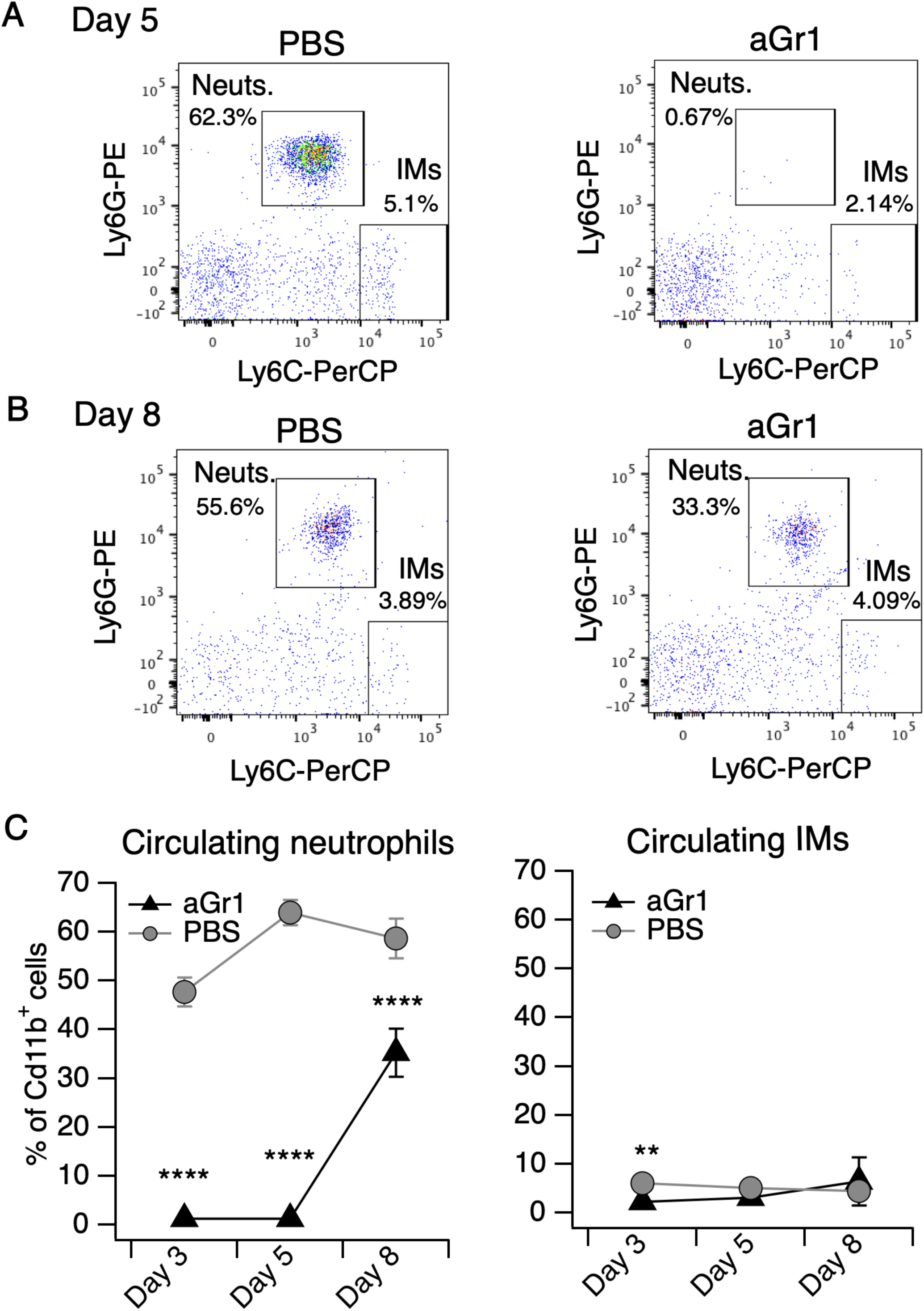
aGr1 treatment preferentially depletes neutrophils. **A.** Representative flow cytometry plots of cells collected from blood of mice injected with PBS or aGr1 (250 µg, i.p.) once-daily for five days concurrent with daily MC903 topical treatment. Shown are CD45.2^+^CD11b^+^ cells, plotted by Ly6G and Ly6C signal, with neutrophil (Neuts.) and inflammatory monocyte (IMs) populations indicated. Neutrophils were defined as Cd11b^+^Ly6G^+^Ly6C^mid/high^ and IMs were defined as Cd11b^+^Ly6G^-^Ly6C^high^ (see Methods). **B.** Representative flow cytometry plot as in **A**, depicting neutrophil and IM populations from blood collected on day 8. **C. (Left)** Neutrophil counts in blood shown as % of Cd11b^+^ cells from aGr1/MC903 (black triangles) and PBS/MC903 (gray circles)-treated animals on days 3, 5, and 8 of the model (two-way repeated measures ANOVA: \*\*\*\**p*_treatment_ < 0.0001, F(1,31) = 299.5; Sidak’s multiple comparisons: \*\*\*\**p_day 3_* < 0.0001; \*\*\*\**p_day 5_* < 0.0001; \*\*\*\**p_day 8_* < 0.0001, n= 16,17 mice). **(Right)** Inflammatory monocyte counts in blood shown as % of Cd11b^+^ cells from aGr1/MC903 and PBS/MC903-treated animals on days 3, 5, and 8 of the model (two-way repeated measures ANOVA: \**p*_treatment_ = 0.0468, F(1,31) = 4.287; Sidak’s multiple comparisons: \*\**p_day 3_* = 0.0015; *p_day 5_* = 0.1918; *p_day 8_* = 0.2013, n= 16,17 mice). Exact values displayed in Figure 2-source data 4.

**Figure 2-Figure Supplement 2.**
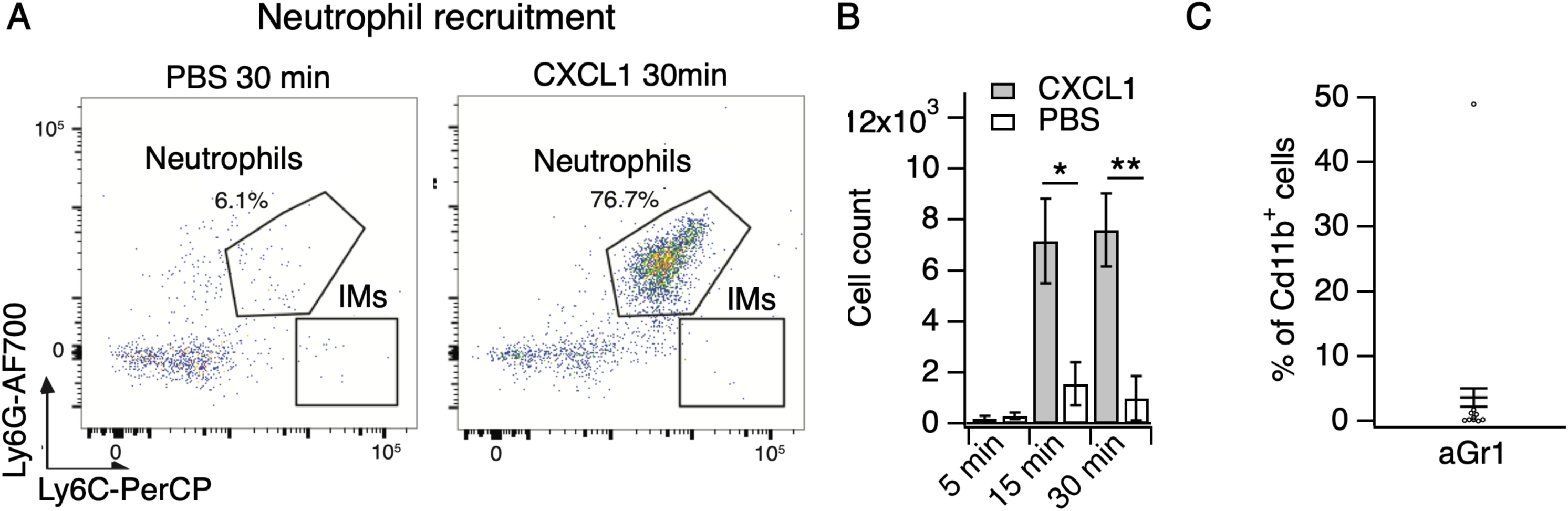
CXCL1 rapidly and selectively recruits neutrophils to skin. **A.** Representative flow cytometry plots of cells from cheek skin of mice injected with PBS or CXCL1 (1 µg in 20 µL, s.c.). Shown are CD45.2^+^CD11b^+^ cells, plotted by Ly6G and Ly6C signal, with neutrophil and inflammatory monocyte (IMs) populations indicated. **B.** Neutrophil count from cheek skin of mice 5, 15, and 30 minutes after injection of CXCL1 or PBS (two-way ANOVA: \**p*_interaction_= 0.0239, F(2,21) = 4.48; Sidak’s multiple comparisons: *p_5 min_* > 0.9999, n=4,5 mice; \**p_day 15 min_* = 0.0141, n=4,4 mice; \*\**p_day 30 min_* = 0.0031, n=3,7 mice). Exact values displayed in **Figure 2-source data 2**. **C.** Blood neutrophils as % of Cd11b^+^ cells approximately 20 hours after injection of 250 µg aGr1 (n=15 mice). Mice assayed for CXCL1-evoked itch behavior immediately preceding blood isolation (see Figure 2E). Exact values displayed in **Figure 2-source data 2**. See Figure 2-Figure Supplement 1C for representative blood neutrophil measurements from PBS-injected animals.

**Figure 2-Figure Supplement 3.**
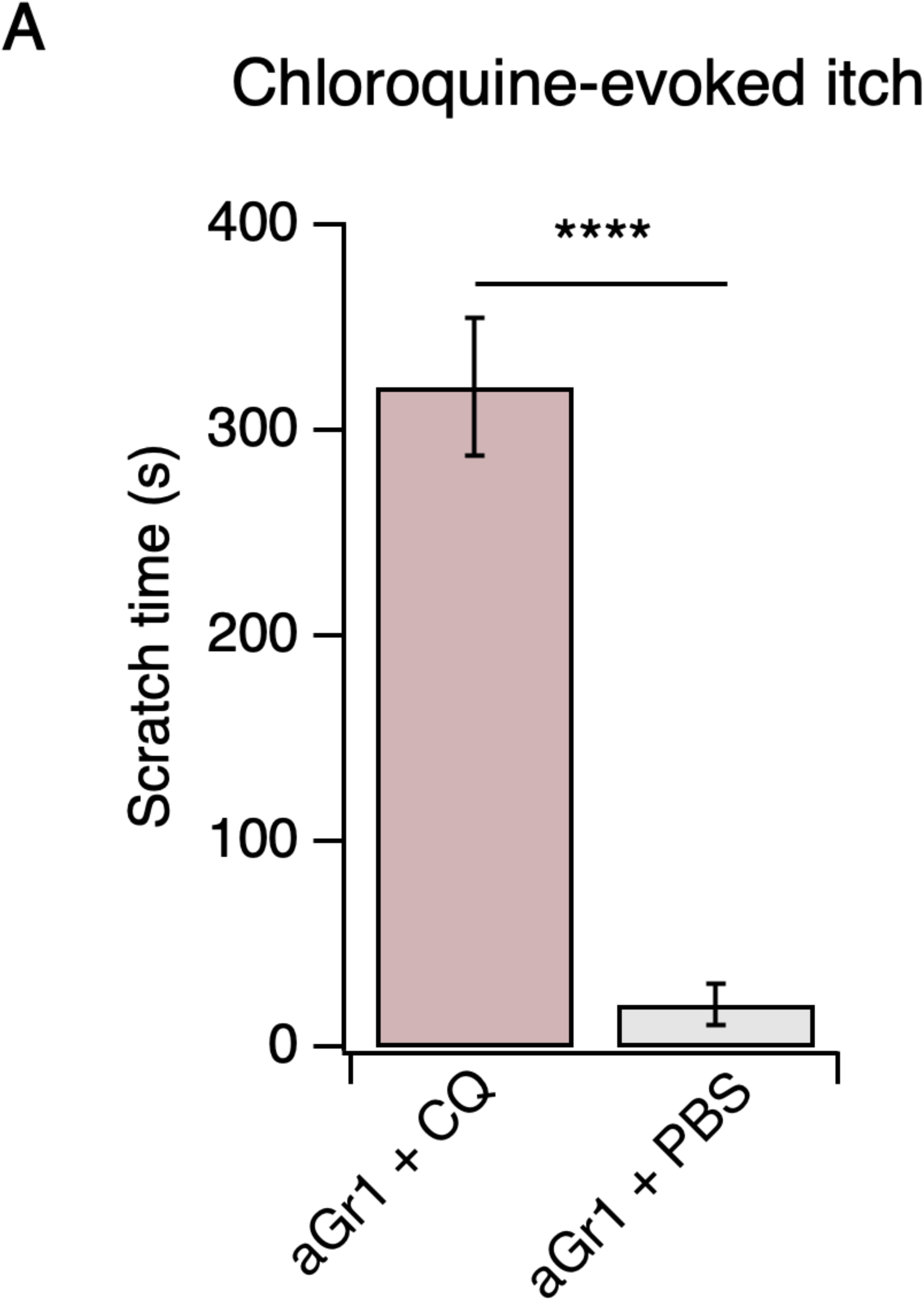
Neutrophil depletion does not affect chloroquine-evoked itch. **A.** Scratching behavior of mice immediately after injection of chloroquine (CQ) or PBS (s.c. cheek). For neutrophil-depletion experiments, mice received 250 µg anti-Gr1 (aGr1) 20 hours prior to cheek injection of CQ or PBS (two-tailed t-test: \*\*\*\**p* < 0.0001 (*t*=10.58, *df*=14); n=6,10 mice). Exact values displayed in **Figure 2-source data 2**.

**Figure 2-Figure Supplement 4.**
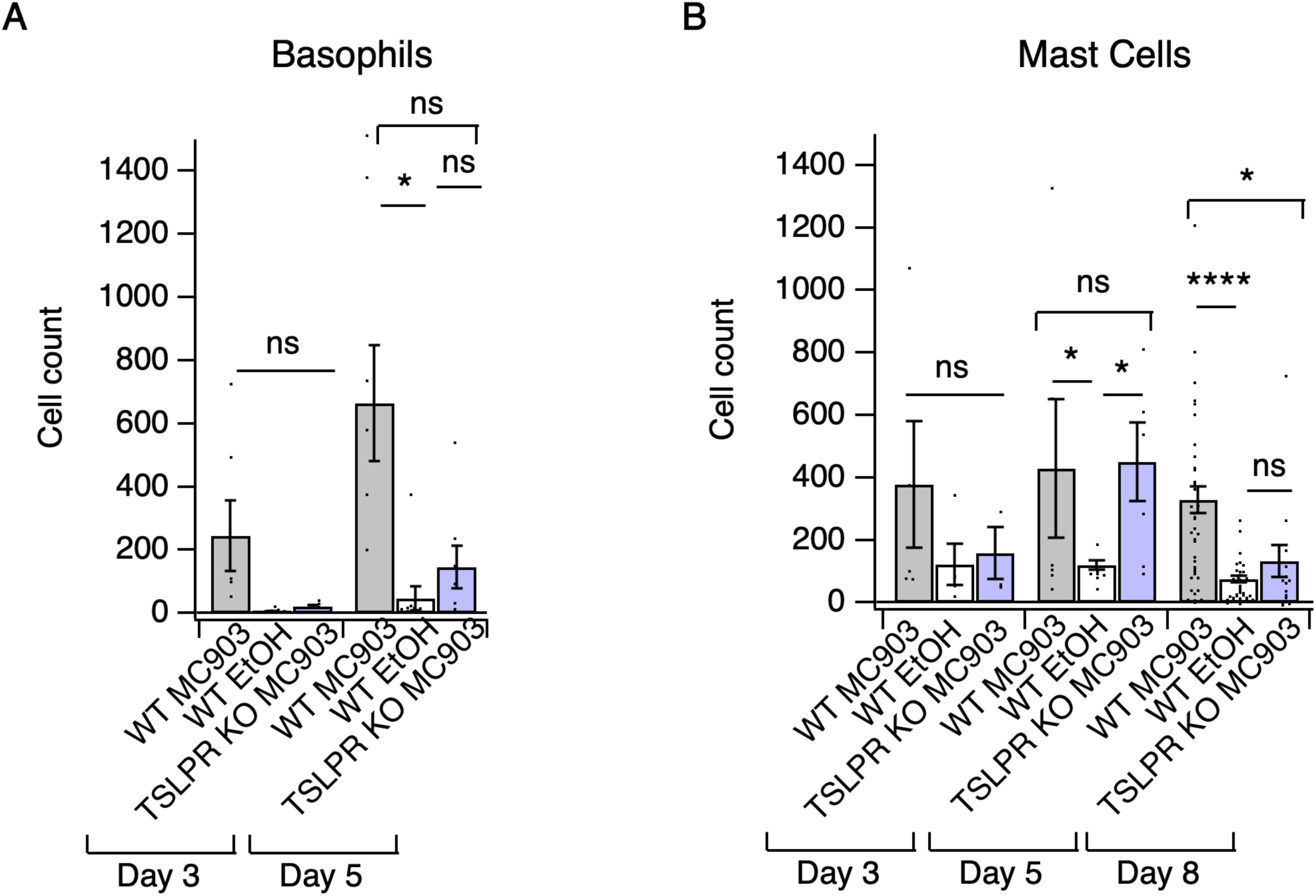
Loss of TSLPR reduces skin basophil and mast cell numbers in the first week of AD development. **A.** Basophil count from cheek skin of wild-type MC903- and ethanol-treated, and TSLPR^-/-^ MC903-treated mice after 3 or 5 days of treatment (two-way ANOVA: \*\*\**p*_time_ = 0.0003, F(2,117) = 8.87; Tukey’s multiple comparisons: *p_day3 WT MC903 vs. KO MC903_* = 0.6540, n=3,5 mice; \**p_day 5 WT MC903 vs. KO MC903_* = 0.1023, n=6,6 mice; *p_day 5 WT EtOH vs. KO MC903_* = 0.9077, n=8,6 mice; *p_day 5 WT MC903 vs. WT EtOH_* = 0.0264, n=6,8 mice). **B.** Mast cell count from cheek skin of wild-type MC903- and ethanol-treated, and TSLPR^-/-^ MC903-treated mice after 3, 5, or 8 days of treatment (two-way ANOVA: \**p*_genotype_ = 0.0384, F(2,117) = 3.35; Tukey’s multiple comparisons: *p_day 3 WT MC903 vs. KO MC903_* = 0.4133, n=3,5 mice; *p_day 5 WT MC903 vs. KO MC903_* = 0.9882, n=6,6 mice; *\*p_day 5 WT MC903 vs. WT EtOH_* = 0.0440, n=6,5 mice; *\*p_day 5 KO MC903 vs. WT EtOH_* = 0.0294, n=6,5 mice; \**p_day 8 WT MC903 vs. KO MC903_* = 0.0188, n=40,15 mice; \*\*\*\**p_day 8 WT MC903 vs. WT EtOH_* < 0.0001, n=40,38 mice; *p_day 8 WT EtOH vs. KO MC903_* = 0.7810, n=38,15 mice). Data from days 3, 5, and 8 are presented in **Figure 2-source data 3.**

**Figure 2-Figure Supplement 5.**
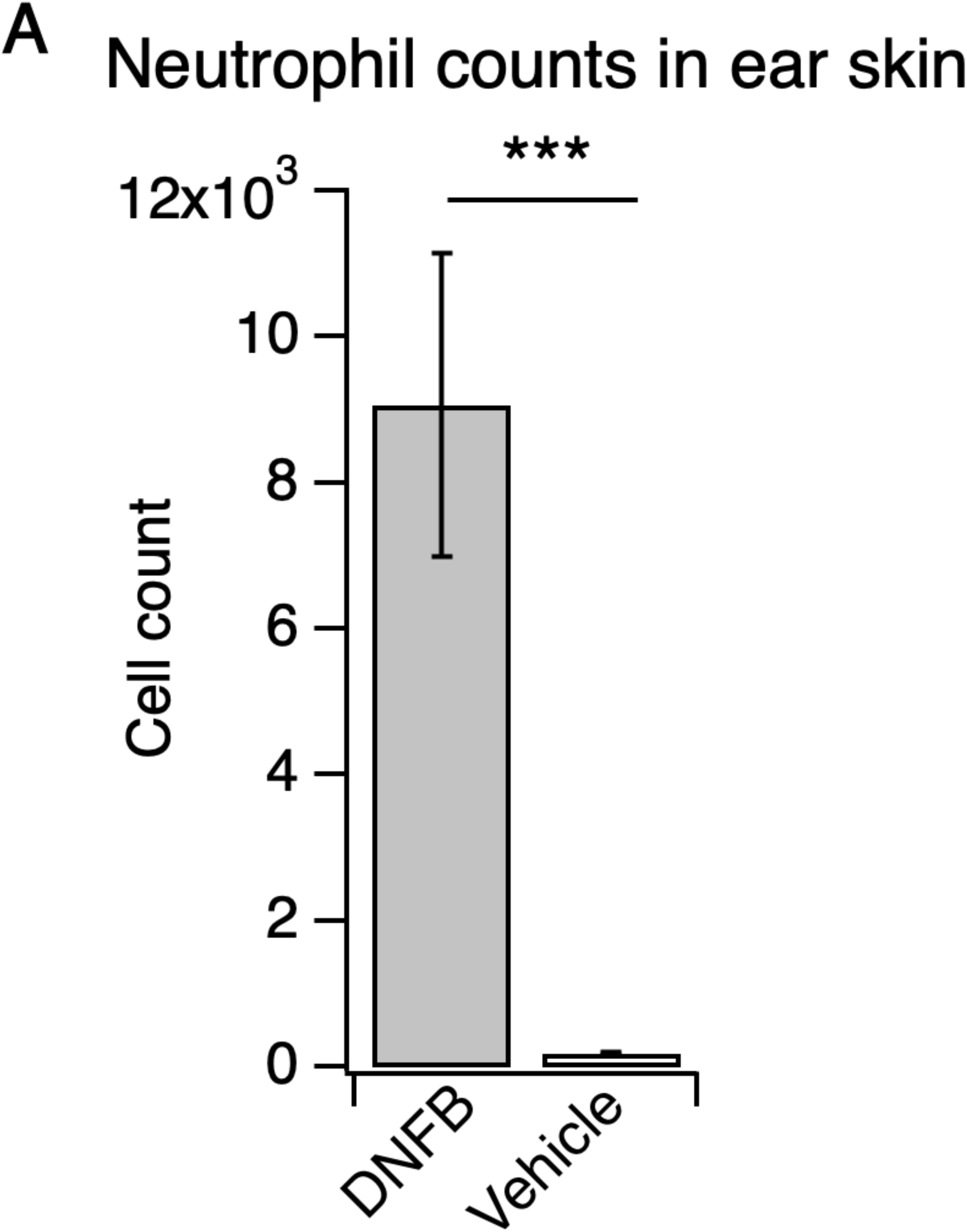
Neutrophils robustly infiltrate the skin in the DNFB mouse model of atopic dermatitis. **A.** Neutrophil count from ear skin of wild-type DNFB- and vehicle-treated mice 24 hours after challenge with DNFB or vehicle performed five days after initial DNFB sensitization on shaved rostral back skin (***p = 0.0004; two-tailed t-test (*t*=4.290; *df* =18); n=10 mice per group). Values from bar plot is reported in **Figure 2-source data 6**.

**Figure 3-Figure Supplement 1.**
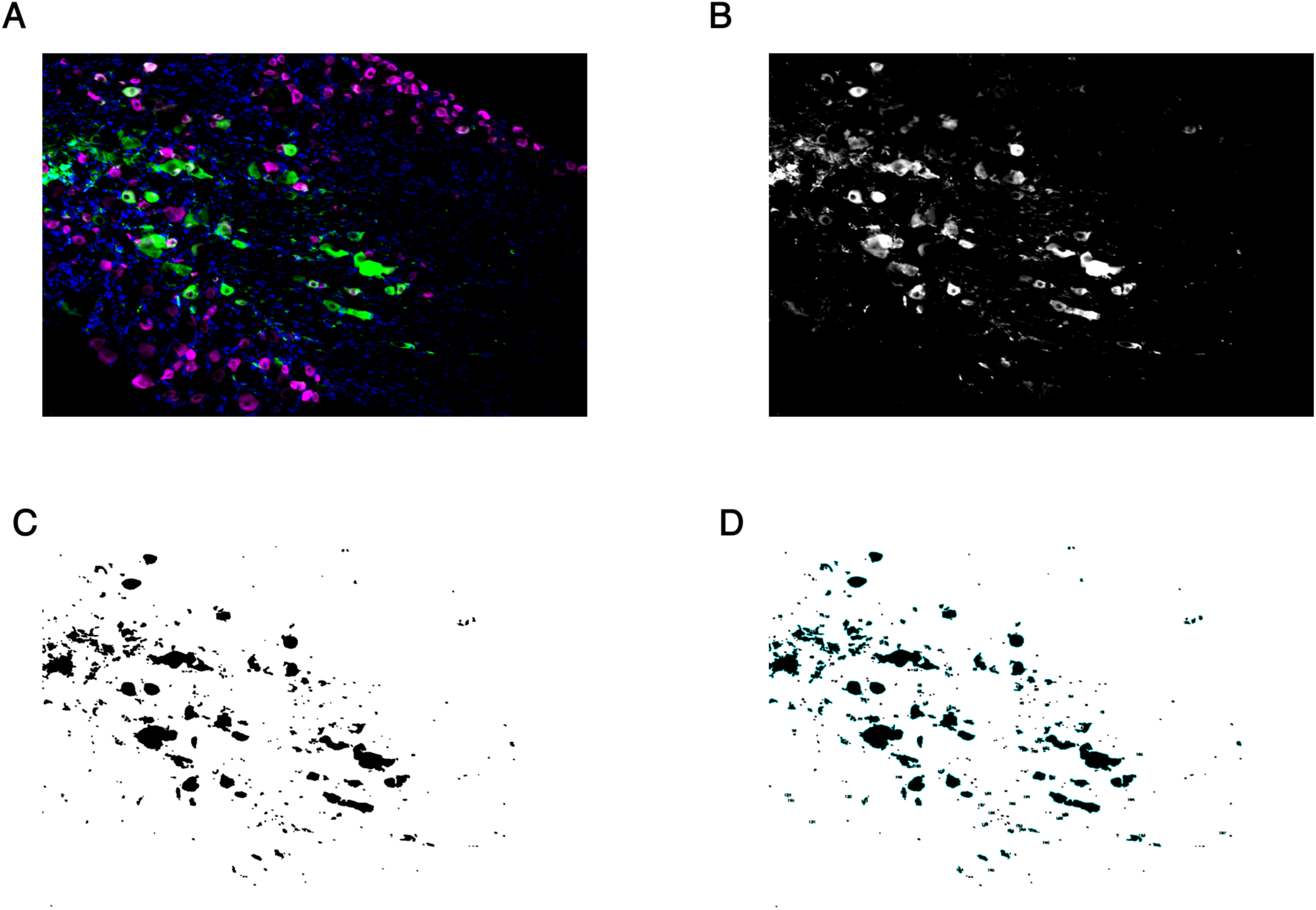
Method of image quantification for sectioned trigeminal ganglia. **A.** Representative composite image showing CD45 (green), Peripherin (magenta), and DAPI (blue). **B.** Single-channel CD45 image with automated min/max intensity thresholding. **C.** Resultant binary image generated from **B. D.** Cells were counted as the number of regions of interest (ROIs) outlined in blue.

**Figure 4-Figure Supplement 1.**
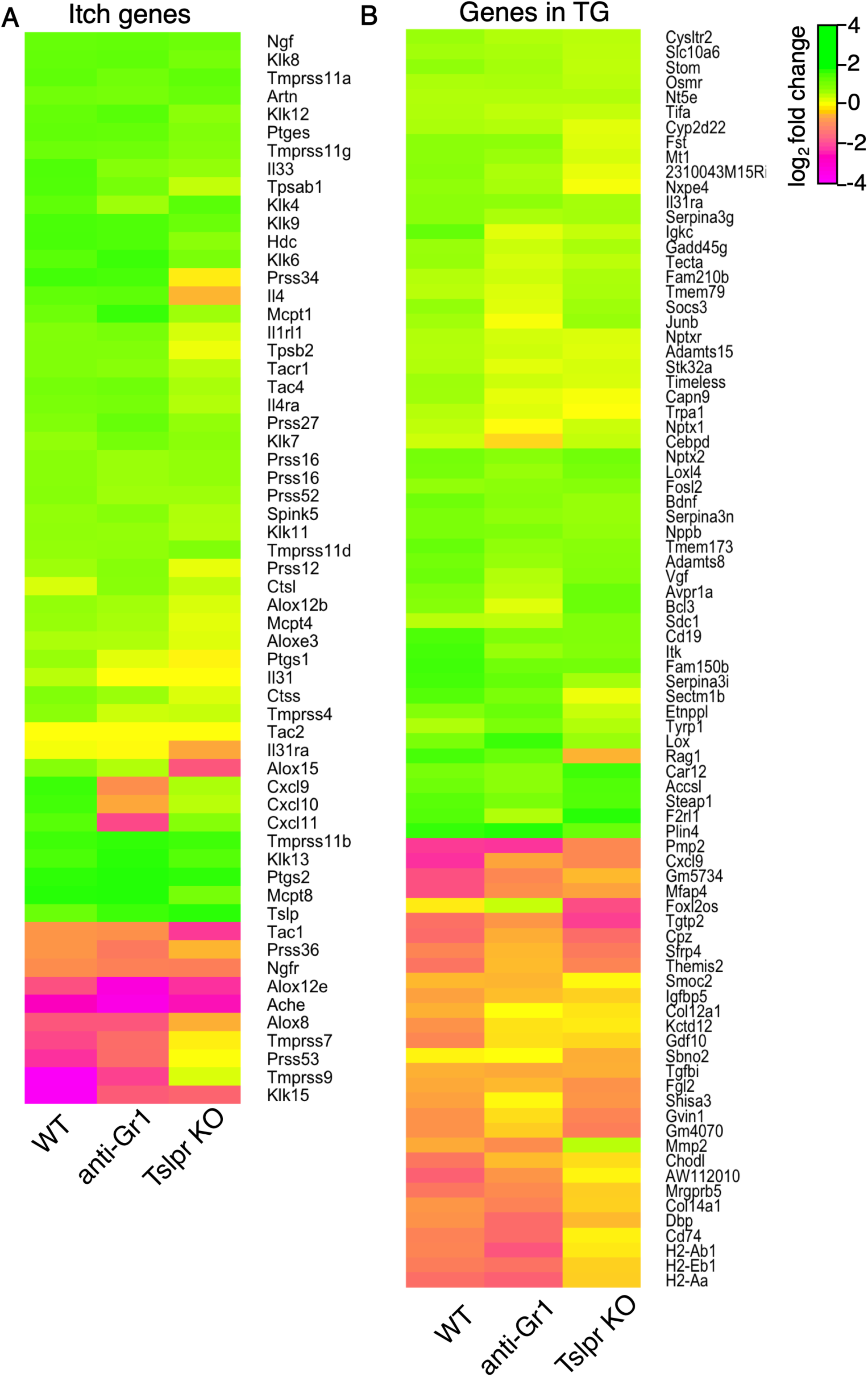
MC903-dependent gene expression changes in aGr1-treated and TSLPR^-/-^ animals. **A.** Heat map showing log_2_ fold change in gene expression (Day 8 MC903 vs. EtOH) for itch-associated genes in wild-type, aGr1-treated, and TSLPR^-/-^ skin. Green bars = increased expression in MC903 relative to ethanol; magenta = decreased expression. Exact values and corrected *p*-values are reported in **Figure 4-source data 3** and **Supplemental Data**, respectively. **B.** Heat map showing log_2_ fold change in gene expression (Day 8 MC903 vs. EtOH) for wild-type, aGr1-treated, and TSLPR^-/-^ mouse trigeminal ganglia (TG) at indicated time points for all genes which were significantly differentially expressed for at least one time point in the MC903 model (See Figure 2D). Green bars = increased expression in MC903 relative to ethanol; magenta = decreased expression. Exact values and corrected *p*-values are reported in **Figure 4-source data 4** and **Supplemental Data**, respectively.

